# Mature parvalbumin interneuron function in prefrontal cortex requires activity during a postnatal sensitive period

**DOI:** 10.1101/2021.03.04.433943

**Authors:** Sarah E. Canetta, Emma S. Holt, Laura J. Benoit, Eric Teboul, Gabriella M. Sahyoun, R. Todd Ogden, Alexander Z. Harris, Christoph Kellendonk

**Affiliations:** Department of Psychiatry, Columbia University Medical Center, New York, NY 10032, USA; Department of Pharmacology, Columbia University Medical Center, New York, NY 10032, USA; Division of Developmental Neuroscience, New York State Psychiatric Institute, New York, NY 10032, USA; Division of Molecular Therapeutics, New York State Psychiatric Institute, New York, NY 10032, USA; Division of Integrative Neuroscience, New York State Psychiatric Institute, New York, NY 10032, USA; Department of Biostatistics, Mailman School of Public Health, Columbia University Medical Center, New York, NY 10032, USA

**Keywords:** Prefrontal cortex, development, parvalbumin, interneurons, gamma oscillations, cognitive flexibility, set-shifting, electrophysiology

## Abstract

In their seminal findings, Hubel and Wiesel identified sensitive periods in which experience can exert lasting effects on adult visual cortical functioning and behavior via transient changes in neuronal activity during development. Whether comparable sensitive periods exist for non-sensory cortices, such as the prefrontal cortex, in which alterations in activity determine adult circuit function and behavior is still an active area of research. Here, we demonstrate that inhibition of prefrontal parvalbumin-expressing interneurons during the juvenile and adolescent period, results in *persistent* impairments in adult prefrontal circuit connectivity, *in vivo* network function and behavioral flexibility that can be reversed by targeted activation of parvalbumin interneurons in adulthood. In contrast, transient suppression of parvalbumin interneuron activity in adulthood produces no lasting effects. These findings identify an activity-dependent sensitive period for prefrontal circuit maturation and highlight how abnormal parvalbumin interneuron activity during development alters adult prefrontal circuit function and cognitive behavior.

## Introduction

In adults, the prefrontal cortex (PFC) is essential for cognitive processes like working memory and behavioral flexibility, and alterations in prefrontal function are believed to underlie impairments in these behaviors in disorders such as schizophrenia and attentional deficit hyperactivity disorder (Brown and Tait, 2016, Millan et al., 2012). The PFC is a late-developing structure, whose circuitry continues to mature through adolescence into young adulthood. Indeed, adolescence appears to be a particularly important time for prefrontal maturation as environmental and pharmacological experiences in this period can elicit long-lasting effects on prefrontal function and behavior (Makinodan et al., 2012, Yamamuro et al., 2018, Bicks et al., 2020b, Thomases et al., 2013, Labouesse et al., 2017). This enhanced sensitivity to environmental risk factors is thought to contribute to the common onset of psychiatric disorders during adolescence (Mayo et al., 2017, Andreasson et al., 1987).

Classic work in the visual system provides an important framework for understanding how transient alterations in environmental experience can lead to persistent effects on adult brain functioning and behavior. In this system, a loss of visual experience during a sensitive period in development, results in activity-dependent remodeling of visual cortical circuitry and long-lasting impairments in visual functioning in adulthood (Hensch, 2005). Recent studies in mice demonstrate that cortical activity may play an important role in sculpting PFC development during an early period spanning late prenatal into early neonatal development in humans (Bitzenhofer et al., 2021). However, extensive refinement of prefrontal circuitry continues through adolescence, and whether neuronal activity during this later period is required for prefrontal circuit maturation remains an open question (Chini and Hanganu-Opatz, 2020).

Within the PFC, parvalbumin-expressing (PV) interneurons regulate the activity of excitatory cortical pyramidal neurons (Markram et al., 2004, Wonders and Anderson, 2006). Moreover, PV interneurons undergo a protracted period of physiological maturation and integration into cortical circuitry during the juvenile and adolescent period, prior to attaining their mature phenotype (Yang et al., 2014, Miyamae et al., 2017). The importance of this cell population for cortical development is underscored by the fact that they show histological abnormalities in multiple neurological and psychiatric disorders with a presumed neurodevelopmental origin, most classically schizophrenia (Lewis, 2014, Ruden et al., 2021). Whether activity of PV interneurons during development is required for proper prefrontal circuit maturation, allowing for optimal cognitive functioning in adulthood, remains unknown.

To address this question, we transiently inhibited PV interneurons in the medial PFC (mPFC) in the mouse during postnatal development using the DREADD receptor, hM4DG_i_ (Roth, 2016), and studied the behavioral and physiological consequences in adulthood. Targeted regions included the prelimbic, infralimbic and cingulate cortex chosen because of their shared functional homology with human dorsolateral prefrontal cortex, which, as previously noted, is implicated in normal cognitive functioning and compromised in psychiatric disorders such as schizophrenia. Using this approach, we found that *transiently* decreasing mPFC PV interneuron activity during juvenile and adolescent development (postnatal day P14-P50), produces *persistent* impairments in adult extra-dimensional set-shifting behavior, with corresponding deficits in PV interneuron-pyramidal cell functional connectivity and task-evoked gamma oscillations. This critical deficit in task-evoked gamma oscillations resulted in a corresponding inability to encode correct vs incorrect task outcome. Additionally, this juvenile period is particularly sensitive for prefrontal circuit maturation as comparable inhibition of mPFC PV interneurons in adulthood did not induce long-lasting changes in behavior and electrophysiology. Strikingly, we could rescue the cognitive behavior and task-induced gamma oscillations in developmentally-inhibited mice by acutely enhancing mPFC PV interneuron excitability in adulthood, demonstrating that cognitive impairments and prefrontal network function can be rescued even in the context of a developmentally altered brain.

Together, this work provides evidence that PV-activity during the juvenile to adolescent time window is required for prefrontal network maturation. These findings highlight how abnormal early activity increases the risk for later maladaptive circuit function and behavior and provide hope that targeted manipulations can reverse developmentally-induced cognitive deficits.

## Results

### A chemogenetic system to transiently inhibit mPFC PV interneurons during development or adulthood

In order to transiently inhibit mPFC PV interneuron activity during discrete windows in development or adulthood we expressed the DREADD receptor, hM4DG_i_, specifically in medial prefrontal PV-expressing interneurons from a young age. Postnatal day 10 (P10) pups expressing Cre recombinase under the PV promoter were injected with an adeno-associated virus (AAV) carrying either a Cre-dependent form of hM4DG_i_-mCherry or the fluorescent marker, mCherry, into the mPFC. Histological assessment of virus expression verified that hM4DG_i_-mCherry was restricted to PV-expressing cells predominantly in the prelimbic, infralimbic and anterior cingulate areas, with some minor spread to the adjacent frontal motor regions of the PFC (Figure 1A and S1). Slice electrophysiological recordings made in hM4DG_i_-mCherry mice at P35 and P115 (Figure 1B, time points chosen to be in the middle of the developmental and adult inhibition windows, respectively) verified the functional effects of hM4DG_i_ in PV interneurons. Bath application of 10 µM clozapine-n-oxide (CNO) hyperpolarized the resting membrane potential of hM4DG_i_-expressing cells at both P35 and P115, while having no effect on mCherry-expressing cells (Figure 1C-D, mean±SEM: −0.1325 mV, n=8 P35 & P115 PV-mCherry cells; −3.99±0.52 mV, 7 P35 PV-hM4DG_i_ cells; −4.88±2.24 mV, 6 P115 PV-hM4DG_i_ cells; 1-way ANOVA, effect of treatment F(2,18)=4.344, p=0.0289; Holm-Sidak post-hoc, P35 PV-hM4DG_i_ v mCherry p=0.035, P115 PV-hM4DG_i_ v mCherry p=0.0297). CNO application also resulted in a rightward shift of firing frequency as a function of input current in cells expressing hM4DG_i_ at both P35 (Figure 1E, n=6 P35 PV-mCherry cells, 6 P35 PV-hM4DG_i_ cells; 2-way ANOVA, effect of input current F(29,540)=87.15 p<0.0001 and effect of treatment F(3,540)=15.43 p<0.0001; Holm-Sidak post-hoc, P35 PV-hM4DG_i_ CNO v P35 PV-hM4DG_i_ ACSF p=0.0013, P35 PV-hM4DG_i_ CNO v PV-mCherry ACSF p<0.0001, P35 PV-hM4DG_i_ CNO v PV-mCherry CNO p<0.0001) and P115 (Figure 1F; n=5 P115 PV-hM4DG_i_ cells, 2-way ANOVA, effect of input current F(19,160)=49.54 p<0.0001 and effect of treatment F(1,160)=10.52 p<0.0014). Cumulatively, these results demonstrate that acute CNO administration decreases the excitability and activity of mPFC PV interneurons expressing hM4DG_i_ both early during development and in adulthood. We also used stereology to quantify the percent of PV cells that were targeted using our viral injection strategy and found that on average 69±7% of PV cells in the prelimbic cortex expressed mCherry and 67±6% expressed hM4DG_i_-mCherry (mean±SEM; n=5 mCherry and n=4 hM4DG_i_-mCherry animals; Figure S1C).

**Figure 1.**
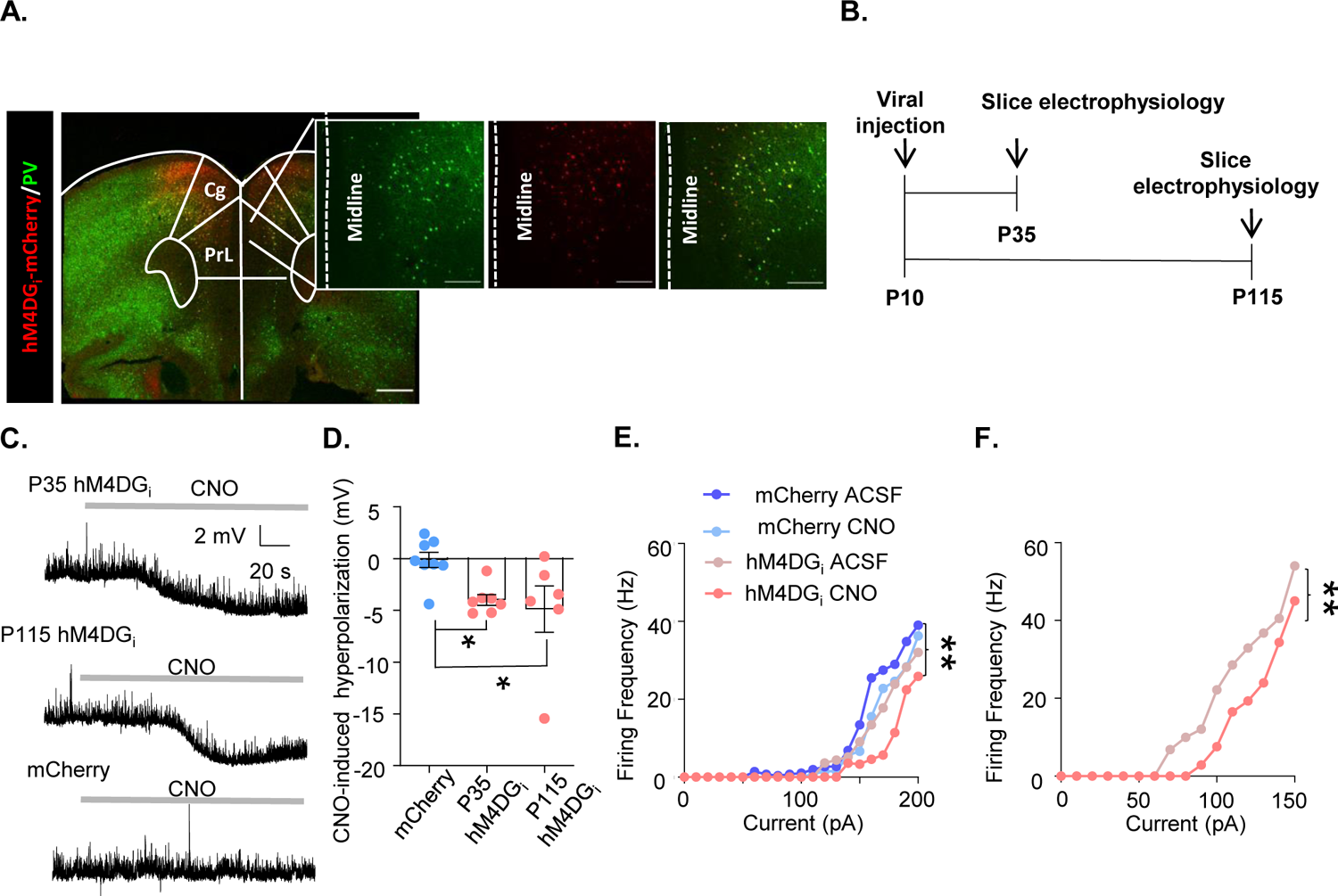
A chemogenetic system to transiently inhibit mPFC PV interneurons during development or adulthood. **(A)** Illustration of hM4DG_i_-mCherry (red) and PV (green) viral expression in prelimbic (PrL) mPFC PV cells. **(B)** Timeline of experiments for validating the function of hM4DG_i_. Mice were injected with virus at P10, and whole cell patch clamp recordings were made at P35 and P115 from cells expressing hM4DG_i_-mCherry or mCherry at baseline and in response to bath application of 10 µM CNO. **(C)** Representative traces illustrating hyperpolarization of the resting membrane potential of hM4DG_i_-expressing, but not mCherry-expressing, cells following bath application of CNO. **(D)** Quantification of CNO-induced hyperpolarization. mCherry expressing cells at P35 and P115 were pooled because CNO did not show an effect at either age. CNO-induced a significant hyperpolarization in P35 and P115 hM4DG_i_-expressing cells relative to mCherry-expressing cells. Dots indicate individual cell responses and bars indicate mean ± SEM. **(E)** Bath application of CNO decreased firing frequency as a function of input current for P35 mPFC PV cells expressing hM4DG_i_, but not mCherry. N=6 cells per group. **(F)** Bath application of CNO also decreased firing frequency as a function of input current for P115 mPFC PV cells expressing hM4DG_i_. N=5 cells per group. Dots and lines in E and F depict mean response. Significance calculated using a one-way ANOVA (D) or two-way ANOVA with post-hoc analysis (E-F). Scale bars are 250 µm. *p<0.05, **p<0.01.

### Developmental inhibition of mPFC PV interneurons results in persistent alterations in behavior and prefrontal network functioning in adulthood

To test whether transiently inhibiting the activity of mPFC PV cells during juvenile and adolescent development results in persistent effects on behavior and prefrontal network function, we injected P10 PV-Cre pups with an AAV expressing hM4DG_i_ or mCherry and administered CNO between P14 and P50 (subsequently referred to as ‘Dev Inhibition’ or ‘Dev Control’, respectively; Figure 2A). We chose P14-50 to test a broad time window that encompasses multiple aspects of mPFC PV interneuron maturation (Miyamae et al., 2017, Yang et al., 2014, Le Magueresse and Monyer, 2013). Forty days following the last injection of CNO, when the animals reached adulthood (≥P90) (Chini and Hanganu-Opatz, 2020), we initiated testing in an odor- and texture-based attentional set-shifting task (the rodent equivalent of the Wisconsin Card Sorting task), which tests cognitive flexibility. In this task, mice dig into two bowls filled with one of two different textures and scented with one of two different odors (Figure 2B). Mice initially learn that odor but not texture predicts reward. During the extradimensional (ED) set-shift the mouse has to learn that the rule shifted from the odor dimension to the texture dimension. We and others have shown that the ED shift is dependent on mPFC PV interneurons (Canetta et al., 2016, Cho et al., 2015, Goodwill et al., 2018, Cho et al., 2020). We also implanted the mice with electrodes to record neural activity in the mPFC during behavior to assess prefrontal network function.

**Figure 2.**
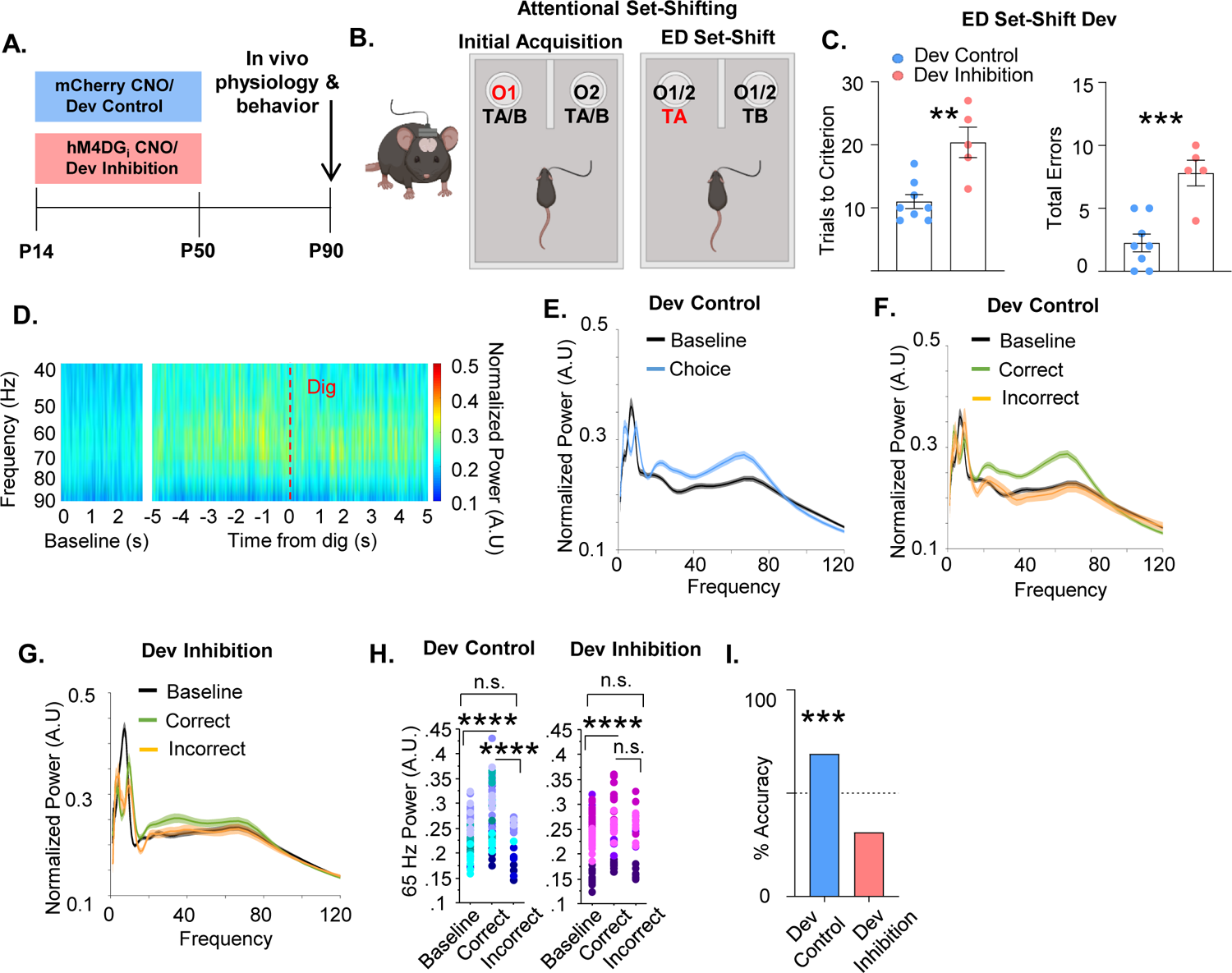
Developmental inhibition of mPFC PV interneurons results in persistent alterations in prefrontal network functioning and behavior in adulthood. **(A)** Experimental timeline. CNO was administered between P14 and P50 to mice expressing hM4DG_i_-mCherry or mCherry in mPFC PV interneurons to produce Dev Inhibition or Control mice, respectively. At P90 mice were evaluated in an attentional set-shifting task while local field potentials were simultaneously recorded in the mPFC as illustrated in the schematic in **(B)**. **(C)** Dev Inhibition mice take significantly more trials to reach criterion (left) and make more errors (right) than Dev Controls. Dots indicate individual animal responses and bars indicate mean±SEM. **(D)** Heat map depicting normalized power (in artificial units, A.U.) as a function of frequency in the first three seconds of the task (baseline) and in a ten second window centered around when the Dev Control animals make their choice (dig, dotted red line). Gamma frequency activity (here shown between 40 and 90 Hz) increases just prior to choice. **(E)** Normalized power versus frequency in the three seconds preceding choice (‘choice’, blue line) versus in the first three seconds of the task (‘baseline’, black line) in Dev Control animals. **(F)** Normalized power versus frequency for the choice-period prior to correct (green) or incorrect (orange) choices, or in the baseline (black) period for Dev Control or **(G)** Dev Inhibition animals. Lines and shading indicate mean ± SEM. **(H)** 65 Hz frequency range power is significantly elevated just prior to correct, but not incorrect, choices relative to baseline for both Dev Control (left, blue dots) and Dev Inhibition (right, pink/purple dots) animals. 65 Hz frequency range power is significantly greater in correct versus incorrect choices in Dev Control, but not Dev Inhibition, mice. **(I)** Choice period 65 Hz frequency range power can predict trial outcome in Dev Control animals (left, blue bar) but not Dev Inhibition animals (right, light red bar). Bar indicates accuracy of model. Dotted line indicates chance level (50%). Significance determined by unpaired t-test (C) mixed-effects linear regression (H), or binomial test (I). **p<0.01; ***p<0.001; ****p<0.0001.

We found that mice that experienced a transient inhibition of mPFC PV interneuron activity during development were impaired in ED set-shifting as adults. Although the developmental inhibition did not affect the initial acquisition (IA) of the task (Figure S2A, mean±SEM: 13.25±1.19 trials, n=8 Dev Control; 13.2±2.01 trials, n=5 Dev Inhibition; unpaired t-test, p=0.9822), it delayed the acquisition of ED set-shifting portion of the task (Figure 2C, mean±SEM: 11±1.12 trials, n=8 Dev Control; 20.4±2.42 trials, n=5 Dev Inhibition; unpaired t-test, **p=0.0021). This impairment was also seen as a significant increase in the total number of errors performed during the ED set-shifting portion of the task (Figure 2C, mean±SEM: 2.25±0.7 errors, n=8 Dev Control; 7.8±1.02, n=5 Dev Inhibition; unpaired t-test, ***p=0.0007). The increased number of errors after developmental inhibition was seen across both perseverative (Figure S2B, mean±SEM: 61.11±15.52%, n=6 Dev Control; 51.39±6.33%, n=5 Dev Inhibition; unpaired t-test, p=0.6042) and random error types (Figure S2C, mean±SEM: 38.89±15.53%, n=6 Dev Control; 48.61±6.33%, n=5 Dev Inhibition; unpaired t-test, p=0.6042). Developmental inhibition did not alter the time it took the mice to complete a trial, suggesting their impairment was not due to a decrease in motivation (Figure S2D&E, IA, mean±SEM: 39.32±15.98 s, n=8 Dev Control; 26.79±7.53 s, n=5 Dev Inhibition mice, unpaired t-test, p=0.5690; ED set-shifting, mean±SEM: 41.76±10.08 s, n=8 Dev Control; 39.95±12.28 s, n=5 Dev Inhibition mice; unpaired t-test, p=0.9123).

Intriguingly, we found that in Dev Control animals, prefrontal power in the gamma frequency increased just prior to when the animals made their choice in the task (signified by digging in one of the pots), relative to gamma power recorded in the beginning of the task (here on referred to as ‘baseline’; Figure 2D-E). Notably, this increase in task-induced gamma was seen only in trials in which the animal proceeded to make the correct choice and not the incorrect choice (Figure 2F). Gamma oscillations can encompass a broad frequency range spanning from 30 to 120 Hz (Sohal, 2016). To capture the oscillation frequency most relevant to the behavior, we identified 62 to 67 Hz (henceforth referred to as 65 Hz) as the frequency range within this 30-120 Hz gamma span where correct trials most frequently showed the maximal difference from incorrect trials (identified using the *max* function in Matlab, Figure S3A) and that showed the largest difference between correct and incorrect trials (Figure S3B). We confirmed that this was the frequency range that was also maximally different between correct and incorrect trials in a second control group of mCherry animals given CNO in adulthood (see Adult Controls below; Figure S3). Correct trials had significantly higher 65 Hz frequency range power relative to incorrect trials and to the baseline period (Figure 2H, 79 baseline/62 correct/17 incorrect trials from 8 Dev control mice, mixed effects linear regression, correct versus baseline p=0.0000, incorrect versus baseline, p=0.462, correct versus incorrect p<0.0001).

Since 65 Hz frequency range power differed between the choice period of correct and incorrect trials, we determined whether it could predict task performance. A machine learning algorithm trained in a subset of 5 correct and 5 incorrect trials using the 65 Hz frequency range window predicted the outcome of remaining trials on a trial-by-trial basis with an accuracy significantly greater than chance (Figure 2I, 69% accuracy, n=54 accurately predicted outcomes of 78 tested trials; binomial test, p=0.0005).

This task-induced increase in 65 Hz frequency range power in the choice period of correct, but not incorrect, trials relative to baseline was also measured in animals that experienced a developmental inhibition of PFC PV interneurons (Figure 2G-H, 69 baseline/41 correct/28 incorrect trials from 4 Dev Inhibition animals, mixed effects linear regression, correct versus baseline p=0.0028, incorrect versus baseline p=0.421). However, in contrast to control mice, 65 Hz frequency range power did not significantly differ between correct and incorrect trials in Dev Inhibition mice (Figure 2H, mixed effects linear regression, correct versus incorrect, p=0.0902). Further, the magnitude of the difference in 65 Hz frequency range power during the choice period of correct versus incorrect trials was significantly higher in Dev Control versus Inhibition mice (mixed effects linear regression, p<0.0001). Finally, 65 Hz frequency range gamma power during the choice period was not able to predict trial outcome with an accuracy significantly greater than chance in Dev Inhibition mice (Figure 2I, 31% accuracy (20/64 trials); binomial test, p=0.999).

These effects are unique to the 65 Hz frequency range. Although the 12-40 Hz frequency range also predicted task outcome in controls (Figure S4, 72% (56/78), p=0.0001), this was also observed in developmental inhibition mice (Figure S4, 66% accuracy (42/64 trials), p=0.008). This suggests that specifically the deficit in task-induced 65 Hz frequency range power may contribute to the impairments in ED set-shifting observed in these mice. No other frequency window predicted task outcome in either controls (Figure S4, 1-4 Hz, 56% (44/78), p=0.154; 4-12 Hz, 51% (40/78), p=0.455; 90-120 Hz, 28% (22/78), p=0.999; 120-200 Hz, 57% (45/78), p=0.106) or developmental inhibition mice (Figure S4, 1-4 Hz, 41% (26/64), p=0.948; 4-12 Hz, 55% (35/64), p=0.266; 90-120 Hz, 42% (27/64), p=0.916; 120-200 Hz, 36% (23/64), p=0.992).

### Adult inhibition of mPFC PV interneurons does not lead to persistent alterations in behavior and prefrontal network function

To determine whether P14-50 is a sensitive time window we inhibited mPFC PV interneurons for a comparable length of time in mature mice (P94-P130, referred to as ‘Adult Inhibition and ‘Adult Control’, respectively; Figure 3A). We choose P94-130 because after P90 traditional markers of PFC maturation have stabilized, including the maturation of PV interneurons (Chini and Hanganu-Opatz, 2020, Delevich et al., 2018, Miyamae et al., 2017, Yang et al., 2014). Adult Inhibition and Control mice took a similar number of trials to reach criteria in both the initial acquisition (Figure S5A, mean±SEM: 11±1.18 trials, n=5 Adult Control mice; 10.4±0.68 trials, n=5 Adult Inhibition mice; unpaired t-test, p=0.6716) and ED set-shifting portion of the task (Figure 3C left; mean±SEM: 10.4±0.93 trials, n=5, Adult Control mice: 12.2±1.59 trials, n=5 Adult Inhibition mice; unpaired t-test, p=0.3576). They also performed a comparable number of errors during ED set-shifting (Figure 3C right, mean±SEM: 2.4±0.93 errors, n=5 Adult Control mice; 3±0.89 errors, n=5 Adult Inhibition mice; unpaired t-test, p=0.6539). There were no differences in the distribution of error types (perseverative or random, Figure S5B-C; mean±SEM: 90±10% perseverative, n=5 Adult Control mice; 66.67±14.3% perseverative, n=5 Adult Inhibition mice; unpaired t-test, p=0.2179; 10±10% random, n=5 Adult Control mice; 33.4±14.3% random, n=5 Adult Inhibition mice; unpaired t-test, p=0.2175) and the latency to complete IA trials (Figure S5D; mean±SEM: 106±29.56 s, n=5 Adult Control mice; 91.64±17.65 s, n=5 Adult Inhibition mice; unpaired t-test, p=0.688) and ED set-shifting trials (Figure S5E; mean±SEM: 47.54±10.29 s, n=5 Adult Control mice; 109.2±53.32 s, n=5 Adult Inhibition mice; unpaired t-test, p=0.2887) was equivalent between the groups.

**Figure 3.**
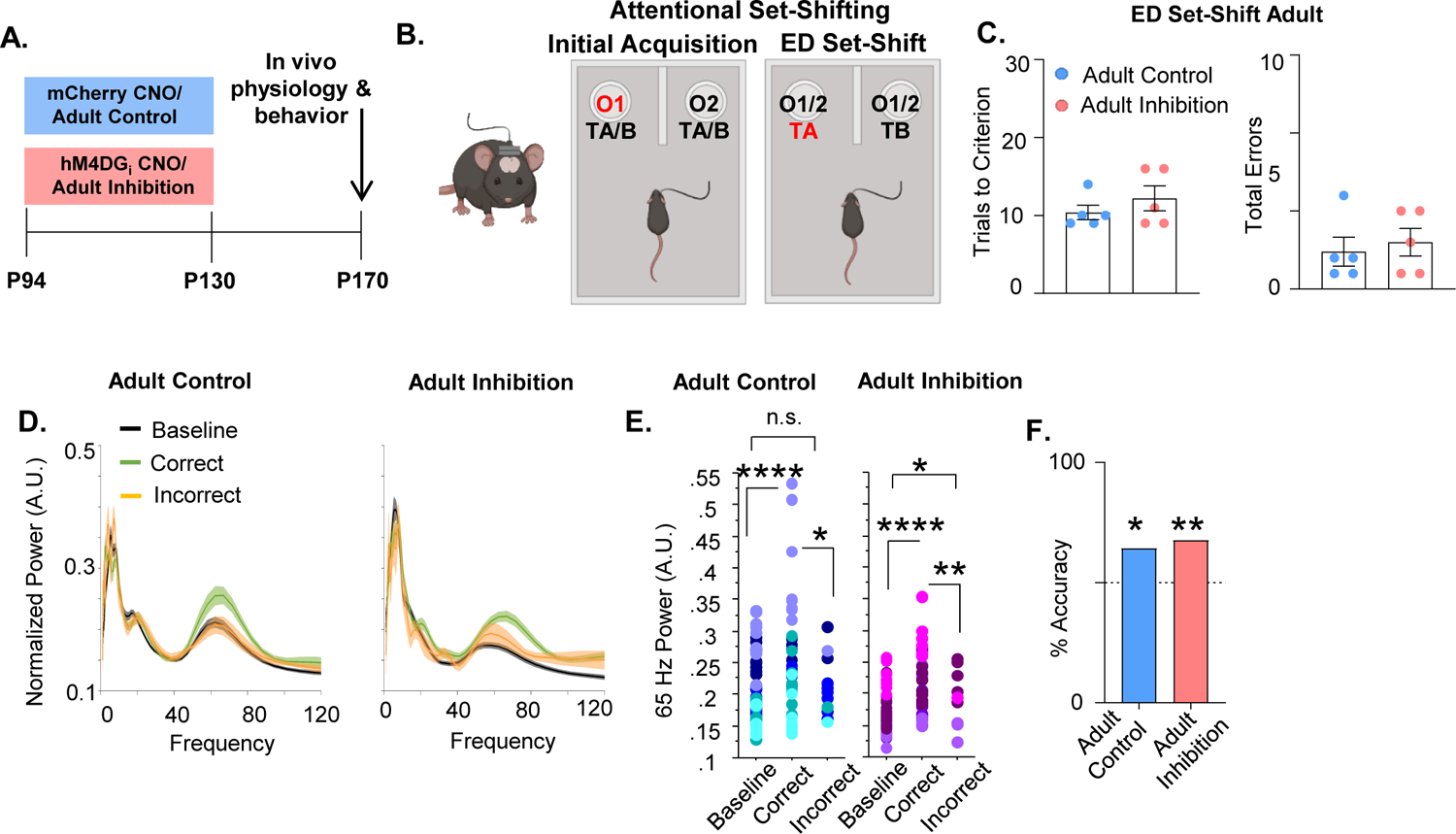
Adult inhibition of mPFC PV interneurons does not result in persistent alterations in prefrontal network functioning and behavior in adulthood. **(A)** CNO was administered between P94 and P130 to mice expressing hM4DG_i_-mCherry or mCherry in mPFC PV interneurons to produce Adult Inhibition or Control mice, respectively. At P170, mice were evaluated in an attentional set-shifting task while local field potentials were simultaneously recorded in the mPFC as illustrated in the schematic in **(B)**. **(C)** Adult Inhibition and Control mice take a comparable number of trials to reach criterion (left) and make a comparable number of errors (right) during ED set-shifting. Dots indicate individual animal responses and bars indicate mean ± SEM. **(D)** Normalized power (artificial units, A.U.) as a function of frequency in the three seconds prior to when the Adult Control (left) or Adult Inhibition (right) mice make a correct (green) or an incorrect (orange) choice relative to baseline (black). Lines and shading indicate mean ± SEM. **(E)** 65 Hz frequency range power is significantly increased just prior to correct, but not incorrect, choices relative to baseline in both Adult Control mice (left; 51 baseline/39 correct/12 incorrect trials from 5 mice, mixed effects linear regression, correct versus baseline p=0.0000, incorrect versus baseline p=0.4412). 65 Hz frequency range power is significantly increased just prior to both correct and incorrect choices relative to baseline in Adult Inhibition mice (right; 44 baseline/34 correct/10 incorrect trials from 4 mice, mixed effects linear regression, correct versus baseline p=0.0000, incorrect versus baseline, p=0.0429). Choice period 65 Hz frequency range power is significantly elevated in correct versus incorrect trials in both Adult Control and Inhibition mice. **(F)** Choice period 65 Hz frequency range power can predict trial outcome in both Adult Control and Inhibition animals with an accuracy significantly greater than chance. Bars indicate accuracy of the models. Dotted line indicates chance level (50%). Significance evaluated by unpaired t-test (C), mixed-effects linear regression (E) and binomial test (F). *p<0.05, **p<0.01, ****p<0.0001.

To ensure that these findings were not a consequence of decreased viral expression over time, we injected a second cohort of mice with the virus in adulthood and again saw no impact of adult inhibition on ED set-shifting behavior; trials to criterion were equivalent between Adult Inhibition and Control mice (Figure S5H; mean±SEM: 10±0.52 trials, n=12 Adult Control mice; 10.18±0.66 trials, n=11 Adult Inhibition mice; unpaired t-test, p=0.8293) as were total errors (Figure S5I, mean±SEM: 1.75±0.37 errors, n=12 Adult Control mice; 1.81±0.50 errors, n=11 Adult Inhibition mice; unpaired t-test, p=0.9131).

In both Adult Control and Inhibition mice, 65 Hz frequency range power during the choice period significantly differed between correct and incorrect trials (Figure 3D&E; 51 baseline/39 correct/12 incorrect trials from 5 Adult Control mice, mixed effects linear regression, correct versus incorrect p=0.0118; 44 baseline/34 correct/10 incorrect trials from 4 Adult Inhibition mice, mixed effects linear regression, correct versus incorrect, p=0.0015). Moreover, in both Adult Control and Inhibition groups, 65 Hz frequency range power predicted trial outcome on a trial-by-trial basis with an accuracy significantly greater than chance (Figure 3F, Adult Control: 64% (27/42); binomial test, p=0.0442; Adult Inhibition: 68% (23/34); p=0.029).

### Developmental inhibition of mPFC PV interneurons results in persistent reductions in their functional inhibition of glutamatergic pyramidal cells in adulthood

Both ED set-shifting behavior and gamma power depend on PV interneuron function. To determine whether developmental inhibition of PV interneurons impairs their functional integration into prefrontal circuitry, mice expressing channelrhodopsin2 (ChR2) and hM4DG_i_ in PV interneurons and treated with either CNO (‘Dev Inhibition) or Saline (‘Dev Control’) from P14-P50, were used for slice electrophysiology at P90 (Figure 4A-B). The strength of GABAergic transmission from PV interneurons onto layer II/III pyramidal cells was measured by evoking neurotransmitter release from mPFC PV cells using blue light and recording the amplitude of the resulting light-evoked inhibitory post-synaptic currents (Le-IPSCs) in patch-clamped pyramidal cells. Le-IPSC amplitudes were significantly reduced after the developmental inhibition (Figure 4C-D, mean±SEM: 2023±367 pA, n=11 Dev Control cells; 1141±205.3 pA, n=17 Dev Inhibition cells; unpaired t-test, p=0.032). This decrease in PV interneuron-mediated inhibition was also reflected as a significant reduction the frequency of spontaneously occurring inhibitory post-synaptic currents (sIPSCs) after developmental inhibition (Figure 4E-F, mean±SEM: 4.81±0.47 Hz, n=12 Dev Control cells; 3.37±0.34 Hz, n=17 Dev Inhibition cells; unpaired t-test, p=0.0165). In contrast, sIPSC amplitude was unchanged (Figure 4G, mean±SEM: 49.76±5.69 pA, n=12 Dev Control cells; 50.37±2.66 pA, n=17 Dev Inhibition cells; unpaired t-test, p=0.8885). Notably, adult inhibition of prefrontal PV interneurons did not persistently alter either the frequency (Figure S6C, mean±SEM: 2.821±0.954 Hz, n=9 Adult Controls cells; 1.857±0.458 Hz, n=10 Adult Inhibition cells; unpaired t-test, p=0.36) or the amplitude (Figure S6D, mean±SEM: 37.87± 3.63 pA, n=9 Adult Controls cells and 44.83±4.77 pA n=10 Adult Inhibition cells, unpaired t-test, p=0.27) of sIPSCs recorded at P170. Together, these results indicate that functional inhibition from mPFC PV interneurons onto pyramidal cells is compromised in adulthood following *developmental*, but not *adult*, inhibition of mPFC PV interneuron activity. The change in frequency, but not amplitude, further suggests a decrease in the quantity or functionality of presynaptic inputs.

**Figure 4.**
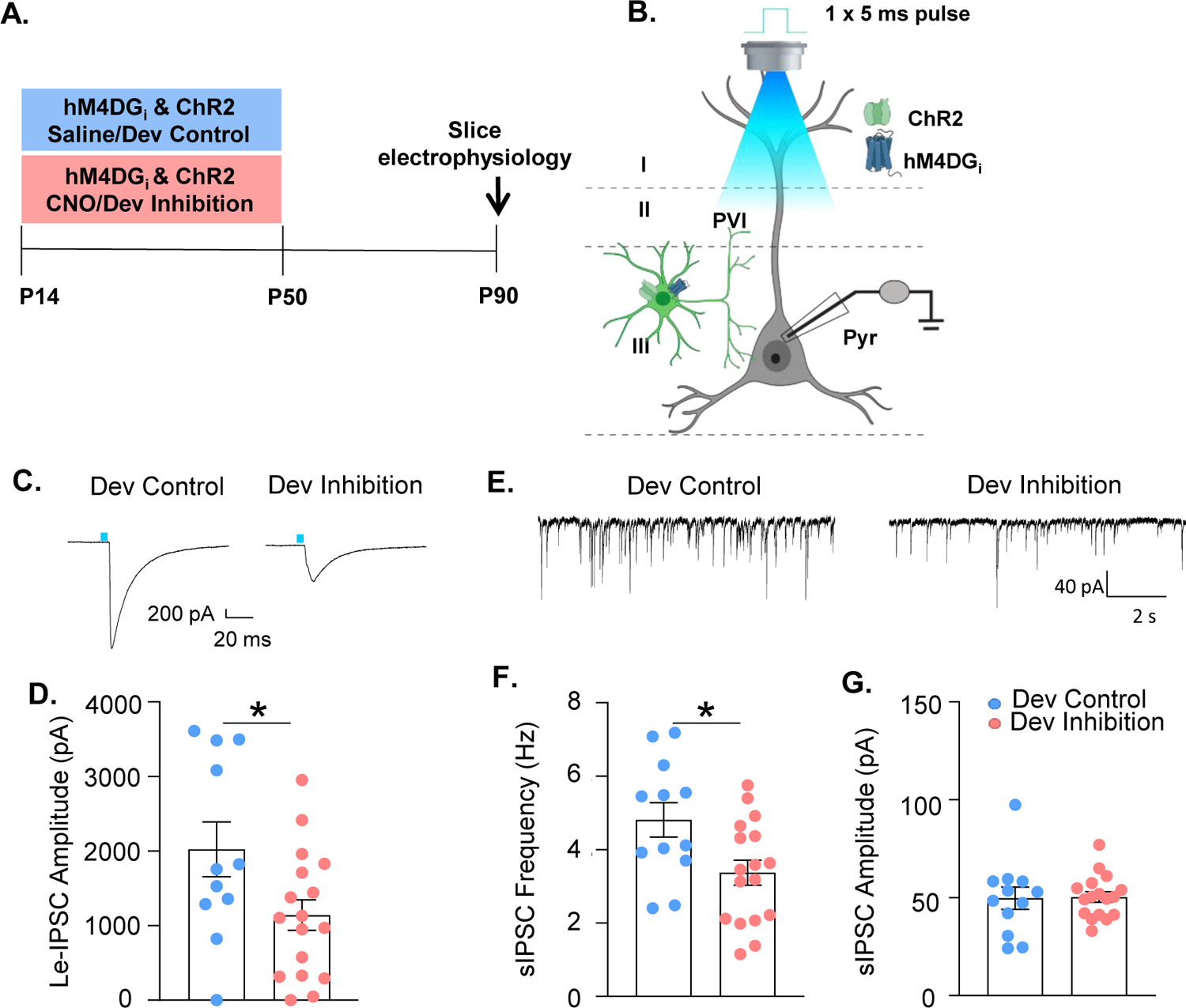
Developmental inhibition of mPFC PV interneurons results in persistent reductions in their functional inhibition of glutamatergic pyramidal cells in adulthood. **(A)** Experimental timeline. Mice expressing ChR2 and hM4DG_i_ in PV cells were administered either CNO (Dev Inhibition) or Saline (Dev Control) from P14-P50. At P90 mice were used for slice electrophysiology. **(B)** Experimental schematic. Whole cell patch clamp recordings were made from pyramidal cells in layer II/III of mPFC from Dev Inhibition and Control mice. The strength of GABAergic transmission from PV cells onto pyramidal cells was measured by evoking neurotransmitter release from mPFC PV cells by delivering a five-ms pulse of 473 nm blue light via the 40x objective and recording the amplitude of the resulting light-evoked inhibitory post-synaptic currents (Le-IPSCs). Spontaneous inhibitory post-synaptic currents (sIPSCs) were also recorded. **(C)** Representative traces showing Le-IPSCs from cells recorded from Dev Inhibition and Control mice. **(D)** Le-IPSC amplitudes were significantly reduced in Dev Inhibition mice. Dots indicate individual cell responses and bars indicate mean ± SEM. **(E)** Representative traces showing sIPSCs from cells recorded from Dev Inhibition and Control mice. **(F)** sIPSC frequency is significantly reduced in Dev Inhibition mice relative to Control mice. Dots indicate individual cell responses and bars indicate mean ± SEM. **(G)** sIPSC amplitude is unchanged. Dots indicate individual cell responses and bars indicate mean ± SEM. Significance evaluated with unpaired t-tests. *p<0.05.

### Developmental inhibition of mPFC PV interneurons results in persistent reductions in the expression of molecular markers

Decreased mPFC PV presynaptic input onto pyramidal cells might result from multiple factors including 1) a loss of PV interneurons, 2) an anatomical reduction of PV interneuron synaptic connections or 3) a physiological change in PV interneurons that renders them less functional. To address the first possibility, the number of PV-expressing cells was stereologically assessed in the mPFC. The number of PV-expressing cells per mm^2^ was not altered by the developmental inhibition (Figure 5F-G; mean±SEM: 5315±1392 cells/mm^2^, n=6 Dev Control mice; 3770±1035 cells/mm^2^, n=7 Dev Inhibition mice; un-paired t-test, p=0.3838). To address the second possibility, we stained tissue for GAD65, which marks presynaptic GABAergic terminals, and assessed the number of GAD65 puncta in close proximity to pyramidal cell soma outlined using the cytoskeletal marker, MAP2 (Figure 5B). We found no differences in the size (Figure 5C; mean±SEM: 2.119±0.49 um^2^, n=7 Dev Control mice; 1.693±0.1 um^2^, n=10 Dev Inhibition mice; un-paired t-test, 0.3312) or number (Figure 5D; mean±SEM: 11.46±1.88 puncta, n=7 Dev Control mice; 10.78±0.74 puncta, n=10 Dev Inhibition mice; un-paired t-test, p=0.7074) of the perisomatic GAD65-expressing puncta. To address the third possibility, we histologically assessed pre-synaptic levels of GAD65 and somatic levels of PV in PV interneurons. Reductions in levels of PV or GAD65 have been associated with decreased functional inhibition (Caballero et al., 2020, Hensch et al., 1998). We then quantified the intensity of PV staining in virus-infected PV interneurons from Dev Inhibition or Control mice. While the intensity of presynaptic GAD65 (Figure 5E; mean±SEM: 107.7±14.76 artificial units (AU), n=7 Dev Control mice; 109.1±9.92 AU, n=10 Dev Inhibition mice; un-paired t-test, p=0.94) and the overall mean staining of somatic PV (Figure 5H; mean±SEM: 59.83±4.12 AU, n=113 cells from 7 Dev Control mice; 58.16±1.88 AU, n=377 cells from 10 Dev Inhibition mice; un-paired t-test, p=0.6844) were not changed, there was a non-statistically significant change in the overall distribution of the PV interneuron staining intensities, with an over-representation of lightly-stained PV interneurons in the Dev Inhibition group (Figure 5I; n=113 cells from 7 Dev Control and 377 cell from 10 Dev Inhibition mice, Kolmogorov-Smirnov test, p=0.0627). As lightly stained PV interneurons have recently been associated with reduced PV plasticity, decreased PV expression may contribute to the deficit in PV functional connectivity (Mukherjee et al., 2019).

**Figure 5.**
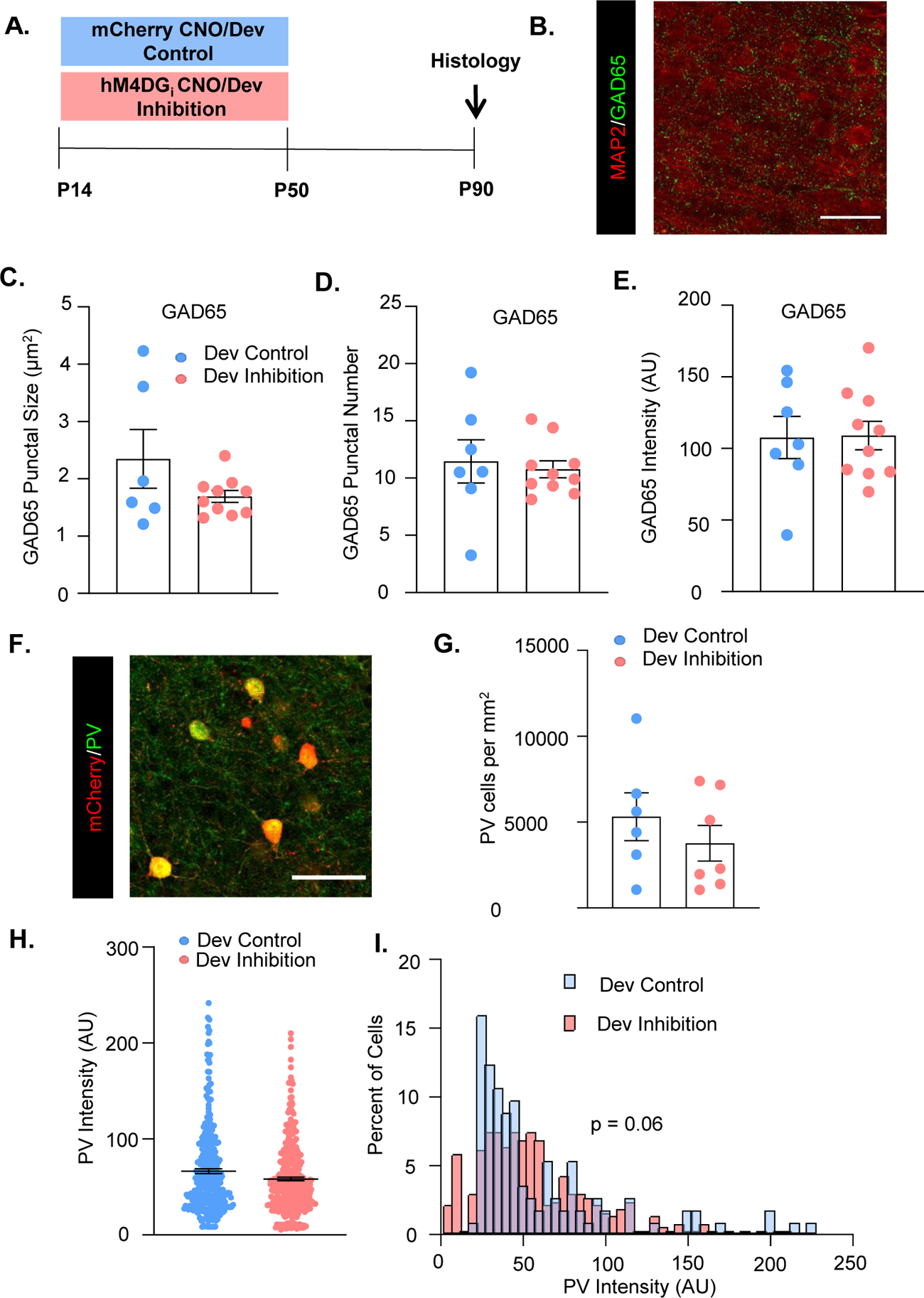
Developmental inhibition of mPFC PV interneurons does not alter PV cell and synapse number but non-statistically significantly decreases somatic PV levels. **(A)** Experimental schematic. P10 PV-Cre mice were injected with either AAV5-hSyn-DIO-hM4DG_i_-mCherry or AAV5-hSyn-DIO-mCherry in the mPFC. CNO was administered between P14 and P50 for the Dev Inhibition and Dev Control mice. At greater than forty days following the end of inhibition, mice were processed for histology. **(B)** Example image of MAP2 (red) and GAD65 (green) staining. **(C)** The size, **(D)** number and **(E)** intensity of GAD65-expressing perisomatic puncta did not differ between Dev Control and Dev Inhibition mice. **(F)** Example image of mCherry (red) and PV (green) staining. **(G)** Stereological counts of PV-expressing cells in the mPFC of Dev Control and Dev Inhibition mice. The number of PV-expressing cells did not differ between these two groups. **(H)** The overall mean intensity of PV in virus-infected cells did not differ between Dev Control and Dev Inhibition mice. **(I)** There was a non-statistically significant difference in the distribution of the intensity of PV expression in virus-infected cells in Dev Control and Dev Inhibition mice. In graphs (C), (D), (E) and (G), dots show individual animal values and bars show the mean ± SEM. In graph (H), dots show individual cell values and bars show the mean ± SEM. Statistical analysis conducted with unpaired t-tests (C-E, G-H) and Kolmogorov-smirnov test (I). Scale bars are 100 µm.

### Enhancing mPFC PV interneuron activity in adulthood with a SSFO rescues the behavioral deficit following developmental inhibition of mPFC PV interneurons

Due to the importance of mPFC PV interneurons for ED attentional set-shifting, we investigated whether we could rescue the deficits in ED set-shifting by acutely and selectively enhancing mPFC PV interneuron excitability in adult mice. To enhance responsiveness of PV interneurons to incoming endogenous activity without imposing an artificial stimulation pattern, we used a stabilized step-function opsin (SSFO) (Yizhar et al., 2011). Dev Inhibition or Control mice also expressing the SSFO in their mPFC PV cells were evaluated in the ED set-shifting task in adulthood. On the first day of testing, half the animals had the SSFO activated by administration of 473 nm light delivered via bilaterally-implanted optic fibers during ED set-shifting, while the other half were in the light OFF condition. Ten days later, ED set-shifting testing was repeated with the SSFO being activated in the animals that had previously been in the light OFF condition, and vice versa (Figure 6A-C). Light activation of mPFC PV interneurons led to a significant reduction in trials to criterion only in Dev Inhibition, but not Dev Control, mice (Figure 6D; mean±SEM: 15.5±1.52 trials light OFF and 10.5±0.85 light ON, n=6 Dev Inhibition mice; 10.2±0.74 trials light OFF and 10±0.84 light ON, n=5 Dev Control mice; 2-way rmANOVA, fixed effect of light F(1, 9)=14.18, p=0.004; no fixed effect of treatment F(1,9)=4.468, p=0.0637; light by treatment interaction F(1,9)=12.08, p=0.007; Bonferroni post-hoc Dev Inhibition light ON v OFF p=0.0009).

**Figure 6.**
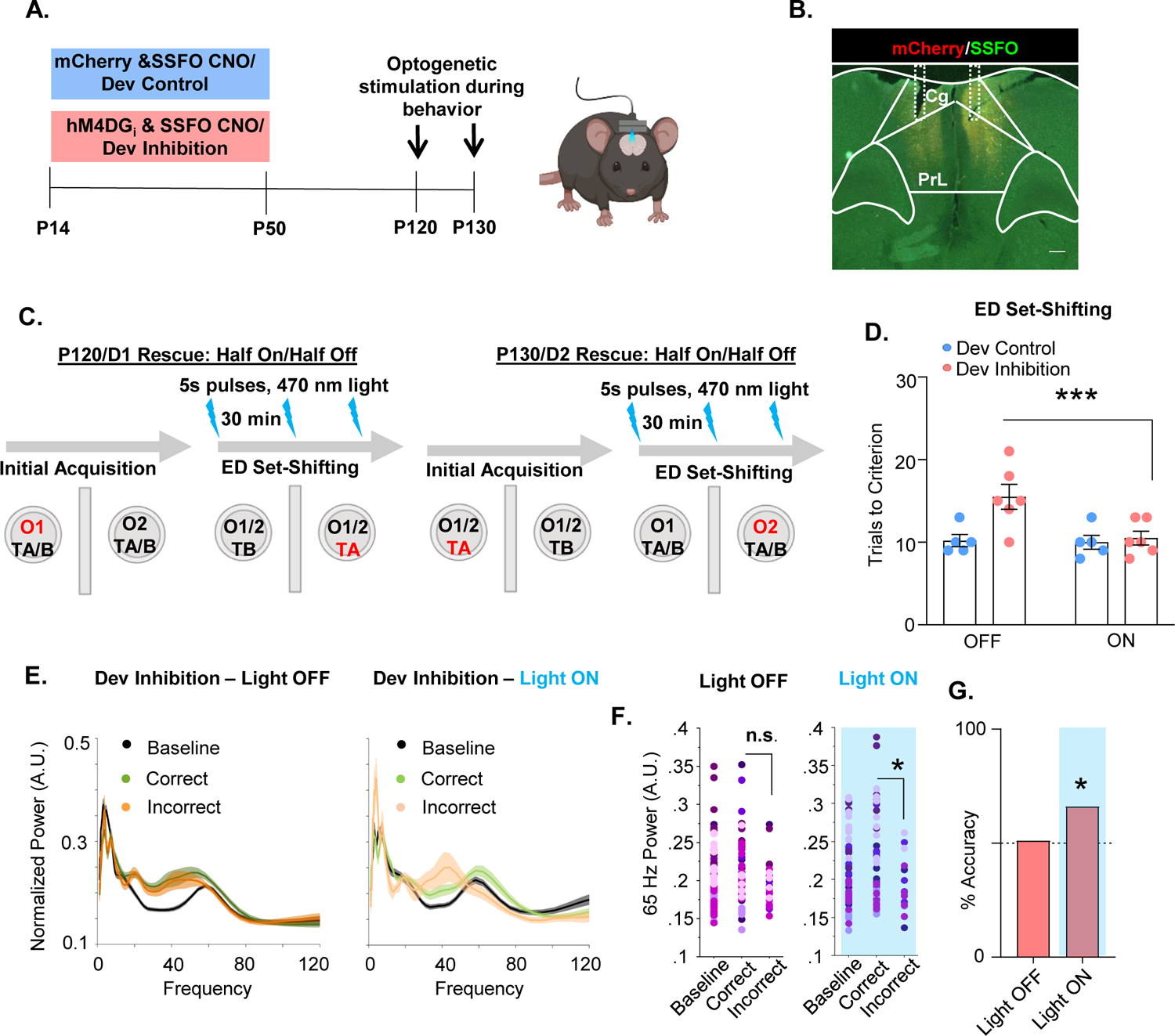
Enhancing mPFC PV interneuron activity in adulthood with a SSFO can rescue behavioral deficits following developmental inhibition of mPFC PV interneurons. **(A)** Experimental timeline. Mice expressing an SSFO in combination with hM4DG_i_-mCherry or mCherry in mPFC PV cells were administered CNO between P14 and P50. In adulthood, optical fibers were bilaterally implanted over the mPFC and mice were evaluated in an attentional set-shifting task with and without optical activation of their PV interneurons. **(B)** Example histology. hM4DG_i_-mCherry or mCherry (red) and SSFO-EYFP (green) were expressed in PV cells in the mPFC. Dotted lines denote bilateral placement of the fiberoptic implants. **(C)** Schematic illustrating the details of the cross-over experiment. Half the animals had the SSFO activated by administration of 473 nm light via bilaterally implanted optical fibers during ED set-shifting on testing day 1 with the other half of the animals in the light off condition. Ten days later, testing was repeated with those animals that originally were in the light off condition receiving SSFO activation and vice versa. **(D)** Following light activation of mPFC PV cells, there was a significant reduction in the number of trials it took the Dev Inhibition animals to reach criterion in ED set-shifting but no change in Dev Controls. **(E)** Power versus frequency for the choice-period prior to correct (green) or incorrect (orange) choices, or in the baseline (black) period for Dev Inhibition animals when the light was OFF (left) or ON (right). **(F)** Choice period 65 Hz frequency range power is not statistically larger in correct versus incorrect trials in Dev Inhibition animals when the light is OFF (left) but when the light is ON choice period 65 Hz frequency range power is statistically greater in correct versus incorrect trials. **(G)** Choice period 65 Hz frequency range power can predict trial outcome in Dev Inhibition animals with an accuracy significantly greater than chance only when the light is ON, but not when it is OFF. Significance assessed by two-way ANOVA followed by post-hoc comparison (D), mixed effects linear regression (F), binomial test (G). *p<0.05, ***p<0.001. Scale bar is 250 µm.

We then determined whether light stimulation not only rescues the behavior but also restored the changes in 65 Hz frequency range gamma power and its ability to predict task outcome. We found that light stimulation enhanced the difference in choice period 65 Hz frequency range LFP power between correct and incorrect trials in Dev Inhibition mice. At baseline, when the light was OFF, Dev Inhibition mice did not show a significant difference in 65 Hz frequency range power in the period preceding correct versus incorrect choices (Figure 6F; 92 baseline/61 correct/31 incorrect trials from 6 mice; mixed effects linear regression, correct versus incorrect, p=0.0744). However, during PV neuron excitation, the difference between correct and incorrect trials became statistically significant (Figure 6F; 63 baseline/49 correct/14 incorrect trials from 6 Dev Inhibition mice; mixed effects linear regression, correct versus incorrect, p=0.0463). Light stimulation also restored the ability of 65 Hz frequency range power to predict trial outcome (Figure 6G, Dev Inhibition Rescue light OFF: 51% (42/82); binomial test, p=0.456; Dev Inhibition Rescue light ON: 66%, (35/53); p=0.0135). These results show that artificially enhancing mPFC PV interneuron activity is sufficient to rescue ED set-shifting behavior and task-induced 65 Hz frequency range power in Dev Inhibition mice with compromised adult mPFC PV interneuron function.

## Discussion

### Identification of a postnatal sensitive period for mPFC PV interneuron development and behavior

Seminal studies in the visual system first identified sensitive periods in which changes in visual experience result in activity-dependent remodeling of thalamocortical inputs that become hard-wired, leading to long-lasting effects on visual functioning (Hensch, 2005, Antonini and Stryker, 1993, Hubel and Wiesel, 1970). Similar activity-dependent remodeling has been shown for the refinement of inhibitory connections made by PV interneurons within the visual cortex during developmental sensitive periods (Baho and Di Cristo, 2012, Chattopadhyaya et al., 2007, Fu et al., 2012, Wu et al., 2012). A major result of our study is that the juvenile/adolescent period is also an activity-dependent sensitive period for prefrontal circuit development. Specifically, we found that transient inhibition of mPFC PV interneurons has long-lasting effects on their functional connectivity, prefrontal network function and behavior. The window we have identified for these effects spans the juvenile period into adolescence (P14-P50). During this time, multiple aspects of mPFC PV maturation typically occur, including physiological alterations in their intrinsic membrane properties and firing properties, strengthening and refinement of their synaptic contacts within prefrontal circuitry, and increases in expression of activity-dependent proteins like PV itself (Caballero et al., 2020, Goodwill et al., 2018, Miyamae et al., 2017, Yang et al., 2014, Bitzenhofer et al., 2020). The dynamic nature of changes occurring to mPFC PV cells in this time is consistent with the dynamic developmental windows that have been shown to be sensitive windows for other brain systems. Accordingly, inhibiting PV interneurons for a comparable length of time during adulthood did not result in persistent behavioral abnormalities.

Our results indicate that PV interneuron activity during adolescence is crucial for mature prefrontal cortical function. Prior work indicates that stimulating pyramidal neurons in the newly forming prefrontal cortex from P7-P11, the human equivalent of late *in utero/*early neonatal development, can lead to alterations in prefrontal circuit function, working memory and social preference that persist into adolescence (Bitzenhofer et al., 2021). Here, we show that even though the adolescent prefrontal cortex is capable of supporting complex cognitive tasks, *endogenous* PV interneuron activity during this time remains crucial for its continuing development. Moreover, the cellular and synaptic changes underlying the effects of altering activity in these two different windows differ, as the P7-11 inhibition resulted in changes in PV cell number, which were not affected after P14-P50 inhibition. Critically, our work highlights that dysfunction in PV interneurons can be a driver of subsequent alterations in prefrontal circuitry and behavior. In the future, it would be important to identify the minimum window of PV interneuron suppression that produces these long-term behavioral effects.

We found that transiently decreasing the activity of mPFC PV interneurons during adolescence persistently impaired adult PV function and ED attentional set-shifting, consistent with the known dependence of ED set-shifting on mPFC PV activity (Canetta et al., 2016, Cho et al., 2015, Goodwill et al., 2018, Cho et al., 2020). In related findings, Mukherjee et al. found that *increasing* activity of mPFC PV interneurons during a late adolescent window (P60-P70) led to persistent rescue of set-shifting behavior in the 22q11 genetic developmental risk factor mouse model (Mukherjee et al., 2019). However, that study left unclear if this intervention simply rectified a genetically-induced deficit or reflected a general developmental sensitive period. If inhibiting endogenous PV activity during their late adolescent window also elicited long-lasting changes in set-shifting behavior, it would be important to understand if the cellular and synaptic changes underlying that behavioral effect are comparable or disparate to those we see following PV inhibition during the juvenile and early adolescent window. Alterations in PV interneuron activity in either developmental window are associated with changes in expression of PV itself. Mukherjee et al. found that this decrease was associated with a decrease in excitatory inputs to PV cells (Mukherjee et al., 2019), and we have evidence that this phenotype can also be associated with a decrease in the strength of inhibition provided by these cells to neighboring pyramidal cells. Therefore, it is tempting to speculate that while the function of PV interneurons may be compromised following alterations in either window, the specific synaptic processes that are disrupted may differ. This remains to be further explored.

### Identification of a task-induced gamma signal predictive of ED set-shifting performance

How prefrontal PV interneurons influence prefrontal network function to enable ED set-shifting behavior remains an open question. We found that in control mice, the power of 65 Hz frequency range gamma activity in the mPFC increases while animals are preparing to make a choice in an ED set-shifting task, relative to a baseline period when they first enter the task. Importantly, this increase is only seen in trials where they go on to make a correct, rather than an incorrect, choice. Prefrontal gamma frequency power is believed to reflect organization of the local pyramidal cell activity by PV interneurons. Therefore, the observed task-induced increase in prefrontal gamma power suggests that mPFC PV interneurons are normally engaged while the animal is preparing to make its choice in the ED set-shifting task. Interestingly, recent work from Cho et al. using *in vivo* calcium imaging to observe bulk changes in mPFC PV interneuron activity during a comparable ED set-shifting task, suggests that mPFC PV cells are activated in incorrect trials, once the animals *realize* they have made an incorrect choice (Cho et al., 2020). We examined this in our dataset by looking at whether 65 Hz gamma frequency power increased in the period after, relative to the period before, making either correct or incorrect choices (Figure S8). In control animals, we found that 65 Hz frequency range power was significantly greater before versus after making a correct choice (Figure S8A-B). For incorrect trials, 65 Hz frequency range power was larger after versus before making the choice, although the effect did not reach statistical significance (Figure S8A-B). In Dev Inhibition mice, gamma power did not change before versus after making either a correct or incorrect choice (Figure S8C-D). It may be that mPFC PV interneurons transiently increase in activity following performance of an incorrect, but not correct, choice and that this transitory peak is better captured with cell-type selective fiber photometry, while gamma oscillations reflect changes in both PV interneurons as well as other elements of the local prefrontal network. Together, our results and those of Cho et al. provide several ways in which mPFC PV interneurons may be engaged and required for different aspects of set-shifting behavior.

65 Hz frequency range power increased before correct choices in all control and experimental cohorts. However, while in control groups the magnitude of 65 Hz frequency range gamma power significantly differed between correct and incorrect trials during the choice period, this difference was not observed in developmental inhibition mice. This decrease in the contrast between correct and incorrect trials may explain why 65 Hz frequency range power predicts trial outcome in the three control groups but not after developmental inhibition of PV interneurons. Notably, stimulation of PV interneurons in adult Dev Inhibition mice rescued the difference in choice period 65 Hz frequency range power and restored the decoding of task outcome and behavioral performance. Cumulatively, our results suggest that mPFC PV interneurons contribute to ED set-shifting behavior by organizing local pyramidal neuron activity in the gamma frequency at the time when the animals prepare to make their choice. This organization allows the flexible grouping/regrouping of neuronal ensembles representing different task contingencies and stimulus environment, which is necessary for adapting to changing stimulus-outcome associations. However, when PV interneurons are inhibited during postnatal development, their functional maturation is impaired affecting their ability to engage the necessary gamma-frequency activity when the mice are adult thus leading to behavioral impairments.

### Postnatal changes in mPFC PV interneuron activity alters their functional integration into cortical circuitry

We also found that developmental suppression of mPFC PV interneuron activity resulted in persistent reductions in the strength of the functional connections made onto local pyramidal cells. This finding is consistent with recent work showing that knockdown of prefrontal PV beginning in early adolescence persistently impaired the cells’ inhibitory output in young adulthood (Caballero et al., 2020). The loss of inhibitory output following our developmental inhibition was not due to a loss of PV-expressing cells as stereological counts of mPFC PV interneurons were unchanged in Dev Inhibition mice. Similarly, an analysis of GAD65-expressing perisomatic puncta indicates that the number of inhibitory synapses onto pyramidal cells is unchanged. Instead, our histological results identified a non-statistically significant overall decrease in the somatic expression of PV itself. PV is a calcium buffer whose expression both influences, and responds to, changes in PV cell activity. Decreased PV expression impairs calcium buffering, altering the cell’s release probability and impeding their ability to sustain firing at their characteristically high frequencies (Caillard et al., 2000). Moreover, normal increases in PV expression across development correspond with increases in the power and peak frequency of gamma oscillations (Bitzenhofer et al., 2020). Additionally, a shift towards lower PV intensity has been associated with reduced plasticity in those cells (Mukherjee et al., 2019). Finally, a reduction in PV expression has been seen in both animal models of genetic and environmental risk factors relevant to psychiatric disorders such as schizophrenia, as well as in post-mortem observations from patients with schizophrenia (Hashimoto et al., 2003, Behrens et al., 2007, Belforte et al., 2010). Our results support the hypothesis that early changes in PV expression have long-lasting consequences for prefrontal network function.

Our findings comport with prior work in visual cortex demonstrating that alterations in the activity of PV interneurons during a developmental sensitive period can impact their integration into cortical circuitry (Chattopadhyaya et al., 2007, Fu et al., 2012, Baho and Di Cristo, 2012, Wu et al., 2012). However, in those studies, alterations in PV activity were primarily found to affect the stability of perisomatic inhibitory synapses made by those cells. In contrast, we find no evidence that our inhibition of mPFC PV interneuron activity influences the number or size of perisomatic inhibitory synaptic contacts. These differences may be explained by our different methodologies for inhibiting cell activity, the use of *in vivo* versus *in vitro* experiments, the sensitivity of the histological assays used to quantify synaptic contacts or differences between the visual and prefrontal cortex. Despite these potential differences in mechanism, both our work in prefrontal cortex, and prior work in visual cortex, indicates that alterations in activity levels of PV cells during discrete developmental sensitive periods, can lead to long-term consequences for the connectivity that these cells make within cortical circuitry. Given that prefrontal network activity is hypothesized to regulate the developmental refinement of prefrontal circuitry and function more broadly, in future studies it will be important to investigate how our inhibition of mPFC PV activity influences the maturation of excitatory cortical connectivity (Chini and Hanganu-Opatz, 2020, Le Magueresse and Monyer, 2013).

### Activation of mPFC PV interneurons rescues prefrontal network function and ED set-shifting deficits

Acutely enhancing mPFC PV interneuron function in adulthood using a SSFO rescued prefrontal network function and the impaired ED set-shifting behavior induced by developmental inhibition of mPFC PV interneuron activity. Given that our developmental inhibition of mPFC PV interneuron activity could have resulted in changes in other cell populations in different brain regions due to downstream effects of the chronic inhibition, these results underscore the centrality of defects in mPFC PV interneurons for the cognitive sequelae resulting from this developmental inhibition. These results are consistent with work demonstrating that optogenetically driving mPFC PV interneurons at gamma frequency (40 Hz) rescues ED set-shifting deficits in adult mice carrying a genetic risk factor for interneuron dysfunction (Cho et al., 2015).

### Relevance to understanding developmental origins of schizophrenia

Alterations in PFC PV interneurons are a hallmark of schizophrenia, a disorder which is also characterized by alterations in cognitive function such as impairments in ED attentional set-shifting (Lewis, 2014, Brown and Tait, 2016). Although schizophrenia typically emerges in late adolescence or early adulthood, it has a suspected developmental etiology due to a plethora of genetic and early environmental risk factors as well as the presence of attenuated symptoms prior to diagnosis (Lewis and Levitt, 2002). However, timing and mechanisms underlying this developmental vulnerability to disease remain unknown.

Our results indicate that developing prefrontal PV interneurons are particularly susceptible to changes in their activity levels during a juvenile and adolescent window. If a comparable vulnerability exists in humans than these results would suggest that any genetic or environmental risk factor that alters prefrontal PV activity could potentially lead to long lasting disruptions in the function of these cells as well corresponding impairments in cognitive behaviors like attentional set-shifting. Both behavioral-level manipulations, such as social interactions (Bicks et al., 2020a), as well as molecular alterations, such as redox dysregulation (Steullet et al., 2017, Sullivan and O’Donnell, 2012), can affect PV activity. Consequently, our results indicate that the timing of exposure to risk factors like social isolation and oxidative stress may be extremely important in determining their long-term effects on behavior. Conversely, these findings suggest that therapeutic interventions delivered within this juvenile and adolescent window might be particularly important in preventing later behavioral dysfunction in people at high risk for developing disease. Finally, our rescue experiments using targeted activation of prefrontal PV interneurons give hope that mechanistically oriented interventions may be able to improve developmentally-induced behavioral dysfunction in adulthood.

## Acknowledgements

The authors thank Julia Greenwald for her assistance with animal husbandry. This work was supported by the Brain and Behavior Research Foundation (grant number 26089) to SEC and (grant number 27384) to AZH and the National Institute of Mental Health (grant number K01MH107760) to SEC, (grant number F31 MH119691) to LJB, (grant number K08MH109735) to AZH and (grant numbers R21 MH121334 and MH117454) to CK. Some figures were created using BioRender.com.

## Author Contributions

Conceptualization, SEC and CK; Methodology, SEC and CK; Formal Analysis, SEC, ESH, LJB, GMS, AZH and RTO; Investigation, SEC, ESH, ET and LJB; Resources, SEC, AZH and CK; Writing – Original Draft, SEC and CK; Writing – Review and Editing, SEC, ESH, ET, LJB, RTO, AZH and CK; Supervision, SEC, AZH and CK; Funding Acquisition, SEC, AZH and CK.

## Materials and Methods

### Animals

All animal procedures were approved by Columbia University’s Animal Care and Use Committee. C57/bl6 (Jackson Labs, Stock #000664) mice were mated with Parvalbumin-Cre (PV-Cre, Jackson Stock #008069) mice to produce heterozygous PV-Cre mice used for early viral injection experiments. For optogenetic slice electrophysiology experiments, mice heterozygous for PV-Cre and homozygous for Ai32 (Cre-dependent channelrhodopsin2 mice, Jackson Labs, Stock *#*024109) were mated with RC::PDi homozygous mice (Cre-dependent hM4DG_i_, gift of Susan Dymecki). Animals were fed *ad libitum* and reared under normal lighting conditions (12/12 light/dark cycle), unless otherwise noted. Male and female mice were used for behavior, *in vivo* and *in vitro* electrophysiology and histology experiments (no effects of sex were seen).

### Surgery

Between postnatal day 9 and 12 (P9 to P12) young mice were anesthetized with a mixture of 40 mg/kg ketamine and 5 mg/kg xylazine and secured in a stereotax outfitted with a mouse pup adaptor using ear cuffs. Viruses, including AAV5-hSyn-DIO-hM4DG_i_-mCherry (#44362, Addgene, 1.2× 10^13 titer), AAV5-hSyn-DIO-mCherry (#50459, Addgene, 4.8 x 10^12 GC/mL titer) and AAV5-Ef1a-DIO-hChR2(C128S/D156A)-EYFP (4.9×10^12 titer; UNC Vector Core, Chapel Hill, NC, USA), were injected bilaterally, targeting the prelimbic region of the mPFC (AP +0.92, ML +/- 0.13, DV −1.45 from skull at Bregma). 0.2 µl of virus was injected at each site over the course of 2 minutes, followed by a 2-minute wait prior to withdrawing the injection pipette.

A similar procedure was used to inject virus in adult mice. Animals were anesthetized with a mixture of 100 mg/kg of ketamine and 5 mg/kg of xylazine. Virus (0.2 µl per site followed by a 2-minute wait) was injected in the mPFC with slightly modified coordinates (AP +1.8, ML +/- 0.35, DV −2.5 from skull at Bregma). Carprofen (1 mg/ml i.p., Zoetis, Parsippany, NJ, USA) was given for post-surgical analgesia. Implantation of the electrode bundle and/or optical stimulation fibers occurred approximately 1 week prior to behavioral testing, to ensure sufficient time for recovery. Adult animals were anesthetized with a mixture of 100 mg/kg of ketamine and 0.5 mg/kg of xylazine (i.p.), and administered 0.05 ml of 1 mg/ml of dexamethasone (subcutaneously; Henry Schein, Melville, NY, USA) prior to surgery, to reduce brain swelling. Bupivacaine (5 mg/ml; Hospira, Lake Forest, Illinois, USA) was also injected subcutaneously at the injection site as an additional analgesic. The electrode bundle composed of 76 µm tungsten wire for the LFP and 13 µm tungsten wire for the stereotrodes (California Fine Wire, Grover Beach, CA, USA) was implanted unilaterally in the left mPFC (AP +1.8, ML +/- 0.35, DV −2.5 from skull at Bregma). The electrode bundle connected to the microdrive (EIB-16 or EIB-32 narrow; Neuralynx, Bozeman, MT, USA) was fixed to a custom 3D-printed stage. Optic fibers were glued to the stage according to bilateral ML coordinates and the electrode bundle was glued to one of the fibers such that it extended 0.02 mm beyond the tip of the fiber.

### Clozapine-n-oxide

(CNO) was obtained from the NIH and stored at −20 °C. A 0.1 mg/ml working solution of CNO was prepared in 0.9% sterile saline at room temperature and administered at a concentration of 1 mg/kg (intraperitoneally, i.p.) twice daily from postnatal day 14 (P14) until P50 (Dev Inhibition/Control and Dev Inhibition/Control with SSFO) or P94 until P130 (Adult Inhibition/Control). The motivation for the dosing of the CNO was based on prior work from our group using hM4D to acutely inhibit other brain circuits (Parnaudeau et al., 2013, Parnaudeau et al., 2015, Carvalho Poyraz et al., 2016), where we use 2 mg/kg. As the effects of CNO have been estimated to last about 8h (Krashes et al., 2011, Roth, 2016, Zhan et al., 2013) we split this in two injections per day with 1 mg/kg. This protocol has been used by others to chronically inhibit neurons during postnatal development (Kozorovitskiy et al., 2012, Zhan et al., 2013, Roth, 2016). A new stock solution of CNO was prepared every two days and stored at room temperature, protected from light.

### Attentional Set-Shifting

Attentional set-shifting was performed as previously described (Canetta et al., 2016). Mice were food-deprived until they reached 80 to 85 percent of their baseline body weight. They received one day of habituation to the set-shifting enclosure (24” L x 11.5” W x 12” H; 10 minutes foraging for pieces of honey-nut cheerios as a reward). That night they were habituated to digging bowls (terracotta cups, 3” diameter, 0.75” height) filled with the bedding media (corn cob or torn paper, Vitakraft, Amazon) and the reward (honey nut Cheerios), which was buried in the bedding media. Habituation was performed overnight in the home cage. Mice then received training/shaping days, in which they were presented with two pots containing samples of either the unscented corn cob or torn paper bedding media baited with a buried honey nut Cheerio reward. Each day the animal received five trials, which continued until the animal found the buried Cheerio reward hidden in each pot. When the animal was consistently digging in each pot within 30-60 seconds following trial initiation, the next day they proceeded to behavioral testing. The first phase of testing was initial acquisition in which the mouse was presented with two pots containing a compound stimulus. The compound stimulus contained a combination of one of two bedding media (corn cob or torn paper) combined with one of two odors (paprika or cinnamon) and only one dimension of the stimulus (e.g. odor, specifically cinnamon) predicted the location of the buried Cheerio reward. For the first five trials of initial acquisition, the animals were allowed to dig in both pots to aid them in acquiring the rule, but the trials were scored as either correct or incorrect based on the first pot in which they chose to dig. Initial acquisition ended when the mice performed 8 out of 10 consecutive trials correctly. The total number of trials to reach this criterion, the number of errors as well as the average latency per trial was analyzed. Once the mouse reached criteria in the initial acquisition portion of the task, it proceeded to the extradimensional set-shifting portion of the task. This phase of the task was identical to initial acquisition, except that the dimension of the stimulus that predicted the location of the food reward was changed (e.g from odor to bedding medium). When the animal performed 8 of 10 consecutive trials correctly, testing was finished. For all trials, if the animal took longer than ten minutes to make a choice in a given trial, the trial was ended and recorded as an error. During ED set-shifting, errors resulting from the mouse making the choice that would have been rewarded in initial acquisition were called ‘perseverative errors’. All other errors were called ‘random errors’. For optogenetic stimulation experiments, set-shifting was performed twice. The first session it was performed as above (in initial acquisition one odor (cinnamon) was the rewarded stimulus dimension and in set-shifting one bedding medium (torn paper) was the rewarded stimulus dimension). The second session was performed 10 days later but paper bedding was the predictive stimulus in initial acquisition and in extra dimensional set-shifting it switched to odor (paprika). Mice performed one session with light ON and one with light OFF in a counterbalanced fashion. We did not observe any training effect between session 1 and 2 for either hM4D (rm 2-way ANOVA, n=7 mice, main effect of light F(1,11)=6.786, p=0.0245; no main effect of session F(1,11)=0.1616, p=0.6954) or mCherry mice (rm 2-way ANOVA, n=5 mice, no main effect of light F(1,6)=0.0438, p=0.8384; no main effect of session F(1,6)=0.04538, p=0.8384). The night prior to the second session of set-shifting, mice were re-introduced to pots containing unscented versions of both bedding medias baited with Cheerios in their home cage. No additional habituation or training was performed prior to the second set-shifting test.

### In vivo electrophysiology

*In vivo* electrophysiology recordings were performed while the animals were performing the set-shifting task. Field potential signals from the mPFC were referenced against a screw implanted in the anterior portion of the skull above the olfactory bulb. LFPs were amplified, bandpass filtered (1– 1000 Hz) and acquired at 2000 Hz with Lynx 8 programmable amplifiers on a personal computer running Cheetah data acquisition software (Neuralynx). The animal’s position was obtained by overhead video tracking (30 Hz) of a LED affixed to the head stage. TTLs were manually inserted to record the timing of relevant events (e.g. trial start, dig, trial end).

Neuralynx files containing LFP data were imported into Matlab with Neuralynx MATLAB import/export package v 4.10. LFP samples were notch filtered using the MATLAB Chronux package to remove 60 cycle noise (http://chronux.org/; rmlinesmovingwinc.m). Mechanical artifacts were eliminated by removing samples whose voltage were more than 3 standard deviations from the entire signal mean. The cleaned signal was then root-mean-squared. Heat maps of normalized power (in artificial units, A.U.) as a function of frequency and time relative to the dig were constructed using the wavelet transformation package in Matlab (https://www.mathworks.com/help/wavelet/ref/cwt.html). Normalized power as a function of frequency was plotted by averaging the data from these plots in the relevant time windows (e.g. 3 seconds before dig for the choice period and the first 3 seconds of the trial for baseline). Any trials less than 6 seconds long were excluded from the analysis because in these trials the choice and baseline periods would overlap. This resulted in exclusion of 9 trials from Dev Control mice (8 correct, 1 incorrect), 5 trials from Dev Inhibition mice (4 correct, 1 incorrect) and 1 correct trial from Adult Control mice. Also note, only 4 of the 5 Dev Inhibition and Adult Inhibition mice used for behavior had usable data for physiology. 65 Hz frequency range power encompassed 62-67 Hz power. Electrode locations were confirmed to be within the mPFC based on location of electrolytic lesions.

### Optogenetic stimulation

Optical stimulation was provided by a laser emitting blue light (473 nm) at 4 mW connected via optical fibers (200 µm, 0.22 NA) to the light fibers (200 µm, 0.22 NA, average 80% transmittance) implanted in the animals’ heads to activate the stabilized step-function-opsin (SSFO). Mice were randomized to receive light ON or OFF in a counterbalanced fashion on one of the two set-shifting test days. During testing, mice received a 5-second pulse of blue light when they were in the familiar environment just prior to beginning ED set-shifting. Mice then received a 5-second pulse of blue light every thirty minutes to maintain the SSFO channels in an open state until testing was complete. All light pulses were administered in the familiar environment between trials.

### Slice Electrophysiology

Whole-cell current and voltage clamp recordings were performed in layer 2/3 pyramidal cells and fast-spiking parvalbumin-expressing (PV) interneurons in the prelimbic region of the medial prefrontal cortex. Recordings were obtained with a Multiclamp 700B amplifier (Molecular Devices) and digitized using a Digidata 1440A acquisition system (Molecular Devices) with Clampex 10 (Molecular Devices) and analyzed with pClamp 10 (Molecular Devices). Following decapitation, 300 µM slices containing the mPFC were incubated in artificial cerebral spinal fluid (ACSF) containing (in mM) 126 NaCl, 2.5 KCl, 2.0 MgCl2, 1.25 NaH2PO4, 2.0 CaCl2, 26.2 NaHCO3 and 10.0 D-Glucose, bubbled with oxygen, at 32° C for 30 minutes before being returned to room temperature for at least 30 minutes prior to use. During recording, slices were perfused in ACSF (with drugs added as detailed below) at a rate of 5 mL/minute. Electrodes were pulled from 1.5 mm borosilicate-glass pipettes on a P-97 puller (Sutter Instruments). Electrode resistance was typically 3–5 MΩ when filled with internal solution consisting of (in mM): 130 K-Gluconate, 5 NaCl, 10 HEPES, 0.5 EGTA, 2 Mg-ATP, and 0.3 Na-GTP (pH 7.3, 280 mOsm).

*Effects of CNO on intrinsic and active membrane properties.* hM4DG_i_-mCherry or mCherry-infected PV cells in the mPFC were identified by their fluorescence at 40x magnification under infrared and diffusion interference contrast microscopy using an inverted Olympus BX51W1 microscope coupled to a Hamamatsu C8484 camera. Intrinsic and active membrane properties (resting membrane potential, input-output firing frequency curve) were recorded in current clamp using the K-Gluconate intracellular solution detailed above before and after 10 µM CNO was bath applied to the slice.

*Le-IPSC and sIPSC Recordings*. Pyramidal cells were visually identified based on their shape and prominent apical dendrite at 40x magnification under infrared and diffusion interference contrast microscopy using an inverted Olympus BX51W1 microscope coupled to a Hamamatsu C8484 camera. Light-evoked postsynaptic inhibitory currents (Le-IPSCs) and spontaneous inhibitory postsynaptic currents (sIPSCs) were recorded in voltage clamp at a holding potential of −70 mV using a high-chloride intracellular solution containing (in mM): 140 CsCl, 4 NaCl, 1 MgCl_2_, 10 HEPES, 0.05 EGTA, 2 ATP Mg^2+^, and 0.4 GTP Mg^2+^ (pH 7.3, 280 mOsm). 20 µM 6-cyano-7-nitroquinoxaline-2,3-dione disodium salt (CNQX, Tocris Bioscience, Briston, UK) and 50 µM D-(-)-2-amino-5-phosphonopentanoic acid (AP5, Tocris Bioscience) were added to the bath to block glutamatergic currents. The cells were placed in the center of the field of view, held at −70 mV in voltage clamp and the current response evoked by a 5-ms pulse of blue light (473 nm) applied by a light emitting diode (LED, Cool LED, Andover, UK) was recorded. The intensity of the LED was set at 1% of maximum intensity. The current trace was filtered with an eight-pole low-pass Bessel filter and the difference between the baseline and the maximum light-evoked current response was recorded. sIPSCs were assessed from 60 seconds of the current recording at a holding potential of −70 mV filtered with an eight-pole low-pass Bessel filter and detected using MiniAnalysis (Synaptosoft, Fort Lee, NJ, USA). All event data was averaged by cell.

### Histology

Adult mice were deeply anesthetized with 100 mg/kg ketamine and 5 mg/kg xylazine (i.p.). For *in vivo* electrophysiology experiments, electrolytic lesions were induced at each recording site by passing current (50 µA, 30 s) through electrodes prior to perfusion. All animals were perfused with phosphate-buffered saline followed by 4% paraformaldehyde in phosphate-buffered saline. Brains were dissected out and post-fixed in 4% paraformaldehyde overnight before being transferred to 1% PBS for long-term storage. Brains were sectioned serially at 50 µm on a vibratome (Leica, Buffalo Grove, IL, USA). The following primary antibodies were used: Parvalbumin (PV; Sigma, Saint Louis, MO, USA, P3088, 1:2000), glutamate decarboxylase 65 (GAD65; Millipore, Billerica, MA, USA, MAB351, 1:1000), microtubule-associated protein (MAP2; Abcam, ab5392, 1:5000), mCherry (rabbit-anti-dsRed; Takara Bio, Mountainview, CA, USA; 632496, 1:500) or green fluorescent protein (GFP; Abcam, Cambridge, UK, ab13970, 1:1000). Primary antibody incubation was 48 hours at 4°C. Alexa Fluor-conjugated secondary antibodies (Invitrogen, 1:1000) were used for secondary detection. Stereology was used to assess PV cell number, as well as PV and virus co-expression, in the prelimbic region of adult Dev Inhibition and Control offspring using StereoInvestigator software (MBF Biosciences, Williston, VT, USA) on a Zeiss epifluorescence microscope (Carl Zeiss Microscopy, LLC, White Plains, NY, USA). GAD65 perisomatic puncta as well as levels of PV present in hM4DG_i_-mCherry and mCherry-expressing cells in the prelimbic region of the mPFC were estimated from analysis of 10x images acquired on a Leica confocal microscope (Leica Microsystems, Buffalo Grove, IL, USA) of sections stained for either PV and mCherry or GAD65 and MAP2, using Image J software (NIH). During image acquisition and quantification, the investigator was blind to the treatment.

Viral expression was confirmed from mCherry or GFP staining, and locations of recording site lesions or optical fiber placements were confirmed under DAPI. Mice were excluded from the analysis if they did not have detectable viral expression in the prelimbic region of the PFC or if the stereotrodes and/or optrodes were not localized within this structure.

### Analysis

Statistical analysis and graph preparation of all data except the *in vivo* electrophysiology was done with Prism 8 software (Graphpad Software, San Diego, CA). *In vivo* electrophysiology analysis was done with custom scripts in MATLAB (Mathworks, Natick, MA) or R (https://www.r-project.org/) and graph preparation of this data was done in MATLAB or Statview (SAS Institute, Cary, NC, USA). We chose to focus on the 65 Hz frequency range (62-67 Hz) because power in these frequencies maximally distinguished correct from incorrect trials in our Dev Control cohort. Although a number of trials also showed a maximal difference at 58 Hz, we chose to avoid encompassing 60 Hz in our range of assessment. We confirmed that this was the frequency range that was also maximally different between correct and incorrect trials in a second independent control group of mCherry animals given CNO in adulthood (Adult Controls). To analyze differences in 65 Hz frequency range power across animals, trials, and groups, we fit linear mixed models with 65 Hz frequency range power as outcome. The random effect was animal, and fixed effects were trial (baseline, correct, or incorrect) and group (Dev Control, Dev Inhibition, Adult Control, or Adult Inhibition). For these analyses only data from trials where the latency to make a choice was >6 seconds was used to avoid an overlap in the time periods considered the choice period and the baseline period. This resulted in the exclusion of 9 Dev Control trials (8 correct, 1 incorrect)), 5 Dev Inhibition trials (all correct), and 1 Adult Control trial (correct). To analyze 65 Hz frequency range power before versus after making a choice in correct and incorrect trials we looked at the difference in mean 65 frequency range power in the three seconds before and the three seconds after making a choice, split by the trial outcome. We again excluded trials where a choice was made in less than 6 seconds, resulting in the exclusion of one Dev Control trial (incorrect). To be consistent with the methods of Cho et al, we only analyzed the first five trials of ED set-shifting for this analysis (Cho et al., 2020). For predictor analysis, the *fitclinear* command from the MATLAB machine learning toolbox was used. The 65 Hz frequency range power for each trial was used as a predictor and choice for each trial (correct vs incorrect) was transformed into binary vectors. To avoid over representation of correct trials, equal numbers of either correct or incorrect choice trials were randomly selected to train the model (10 total, 5 correct and 5 incorrect). This model was then tested on the remaining trials and the probability of achieving the observed number of successes given a theoretical distribution based on 50% accuracy was determined with a binomial test. For the predictor analysis, all trials were included regardless of the time it took to make a choice since the choice period was not being compared with the baseline period.

## Supplemental Figure Legends

**Figure S1 (related to Figure 1).**
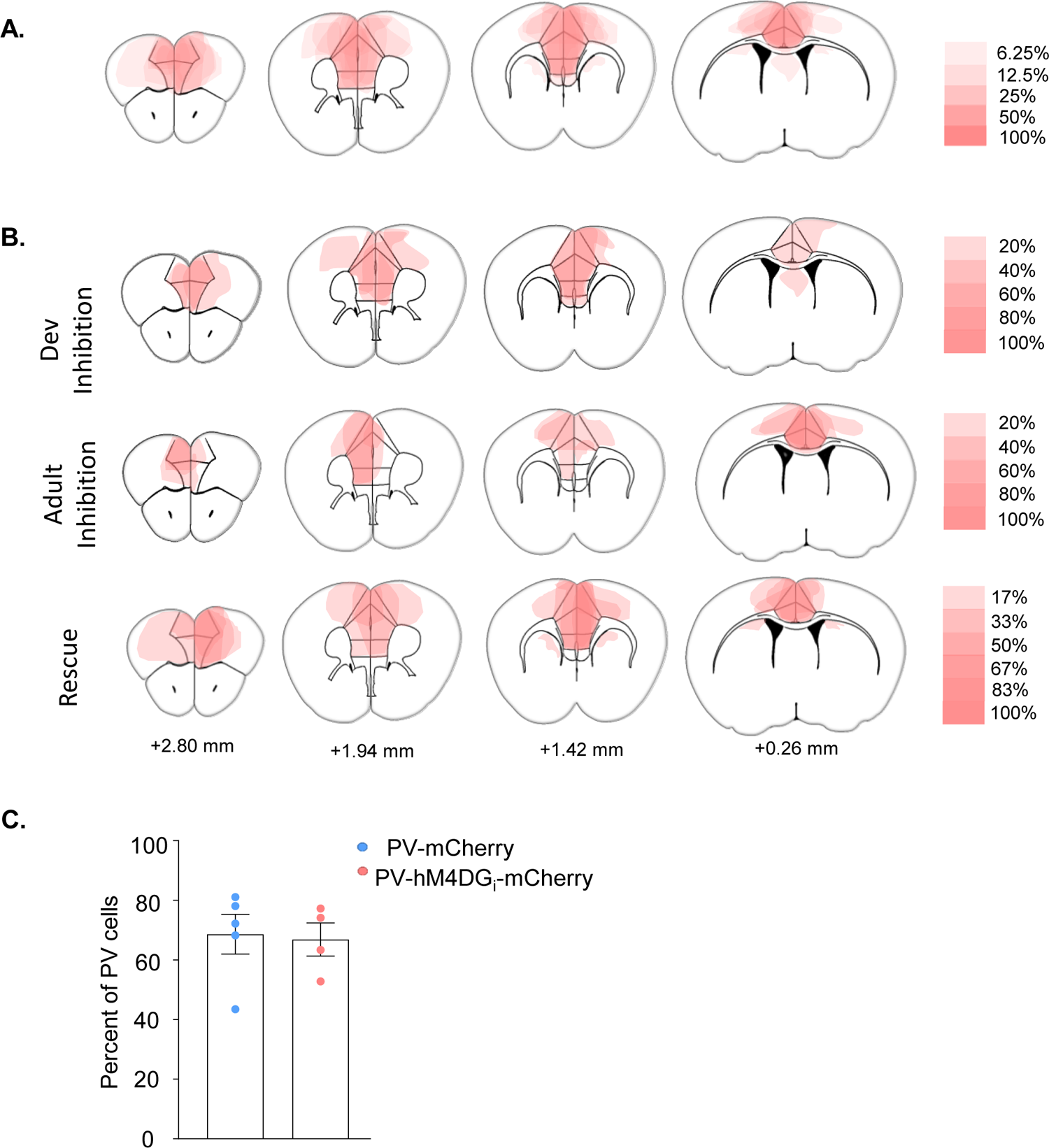
Characterization of viral expression. **(A)** Expression profile of all hM4DG_i_-injected animals (n=16). Heat index depicts percent of animals with expression in a given region. **(B)** Expression profile of hM4DG_i_-injected animals shown separately for developmental inhibition (n=5), adult inhibition (n=5) and rescue (n=6) cohorts. Heat index depicts percent of animals with expression in a given region. **(C)** The percent of PV cells in prelimbic cortex co-expressing either the mCherry or hM4DG_i_-mCherry virus was quantified using stereology. Bars show the mean ± SEM and dots show individual animal values.

**Figure S2 (related to Figure 2).**
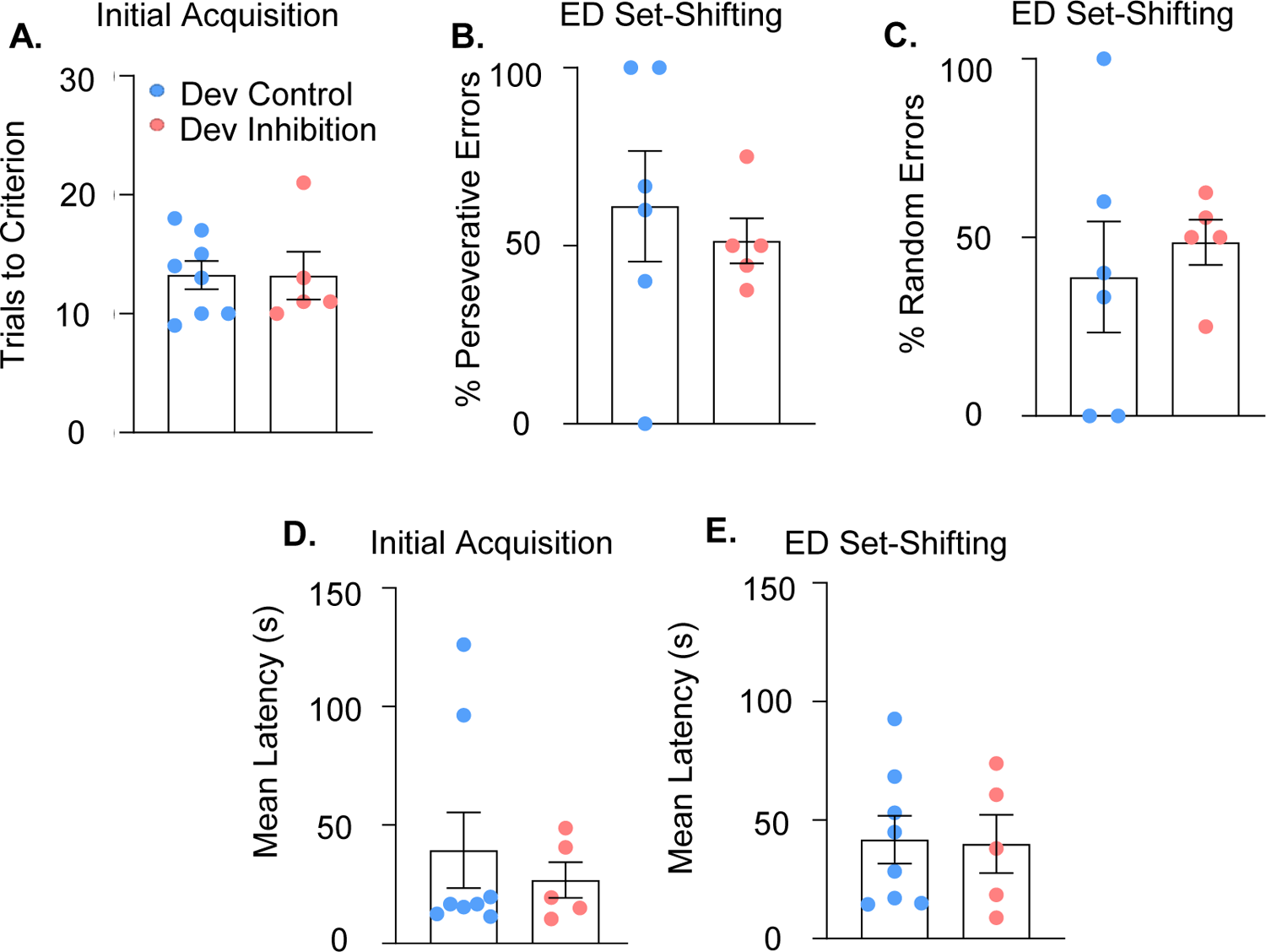
Additional behavioral analysis related to the mPFC PV interneuron developmental inhibition. **(A)** Trials to reach criterion in the initial acquisition phase of the set-shifting task is not affected by developmental inhibition of mPFC PV interneurons. **(B)** There is no difference in the percentage of perseverative or **(C)** random errors made by Dev Inhibition relative to Control mice. Note: 2 Dev Control animals did not make any errors. Dev Inhibition and Control animals have a comparable mean latency to make a choice in both the initial acquisition **(D)** as well as extradimensional phase **(E)** of the set-shifting task. In all graphs, dots show individual animal values and bars show the mean ± SEM. Statistical analysis performed using unpaired t-tests.

**Figure S3 (related to Figure 2).**
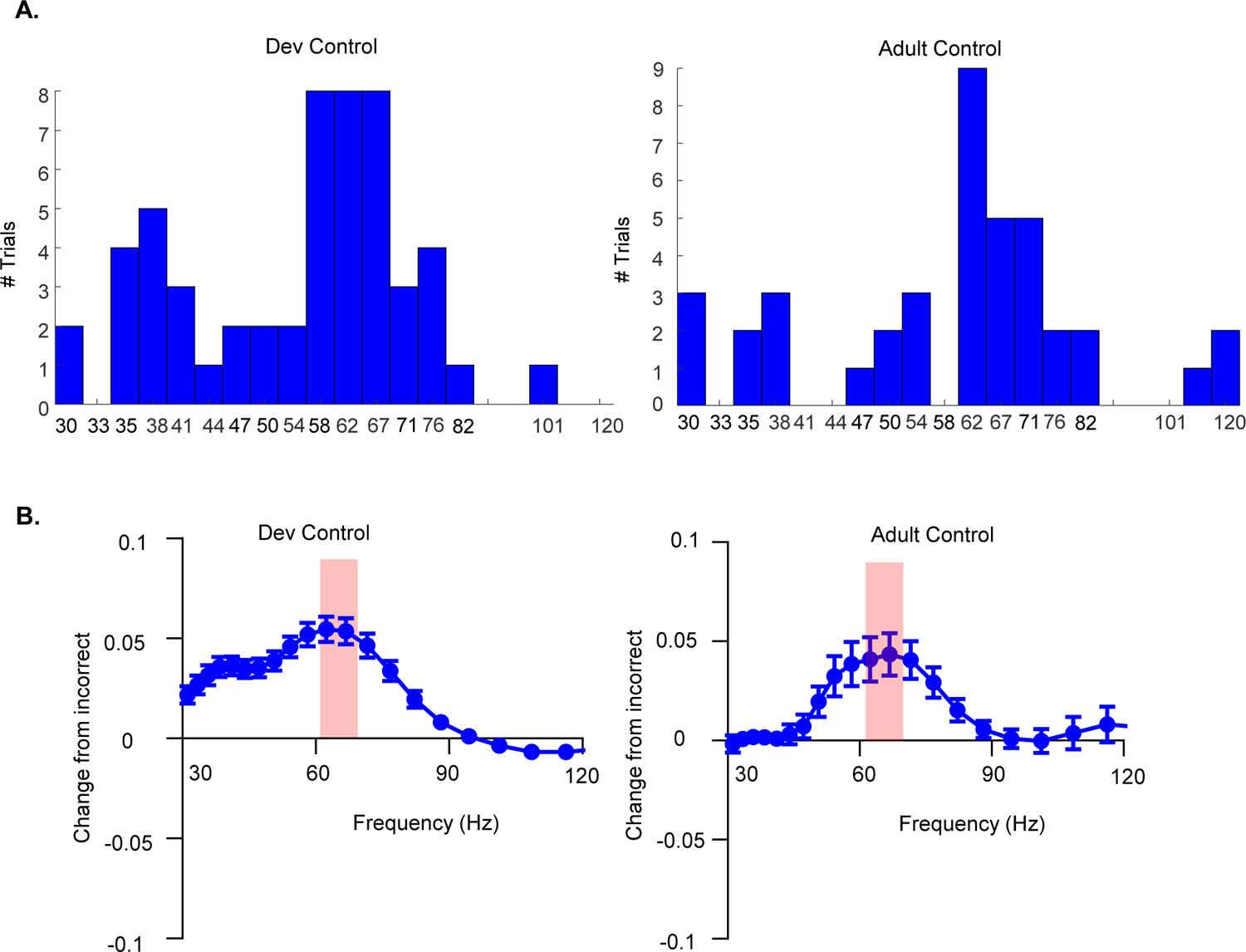
Additional in vivo electrophysiological analysis related to the selection of the frequency range for analysis. **(A)** Histogram for the number of correct trials per frequency for which the difference in power between correct and incorrect trials is larger than the difference in power in all other frequencies for Dev Control mice. Data were not included for the two Dev Control animals that had no incorrect trials. Although a number of trials also showed a maximal difference at 58 Hz, we chose to avoid encompassing 60 Hz in our range of assessment. **(B)** The difference in LFP power across frequencies in the choice period for each correct trial versus the mean power across frequencies for all incorrect trials for a given mouse. Dots show mean values and bars show the SEM. Red shading indicates 62-67 Hz where this correct versus incorrect difference is maximal.

**Figure S4 (related to Figure 2).**
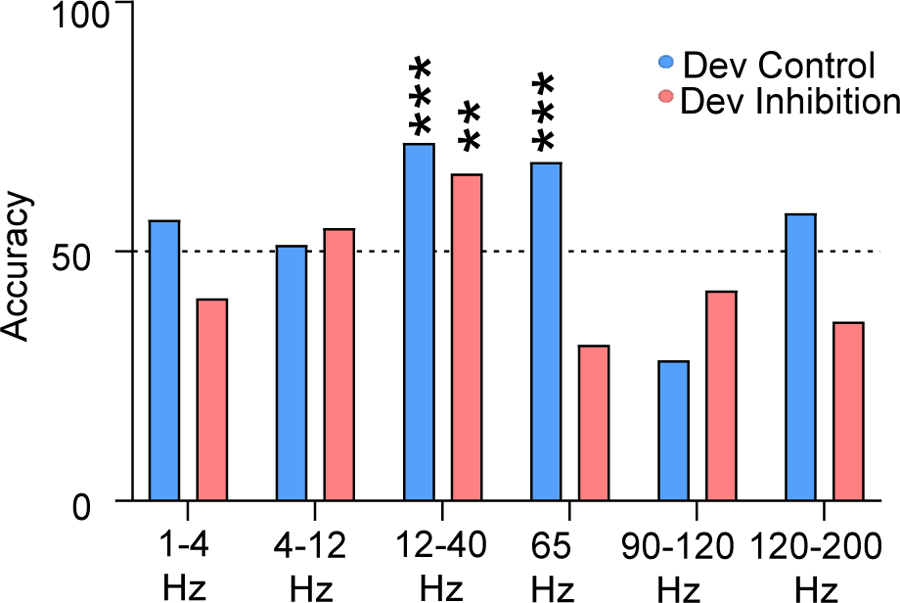
Analysis of the power of different frequency ranges during set-shifting for Dev Inhibition and Control animals. **(A)** 1-4, 4-12, 90-120 and 120-200 Hz mean power cannot predict trial outcome in Dev Control or Dev Inhibition animals with an accuracy significantly greater than chance (blue bars). While 12-40 Hz power can accurately predict outcome for both Dev Control (blue) and Dev Inhibition (light red) mice, 65 Hz frequency range power only accurately predicts power for Dev Control animals. Bars indicate accuracy of the model. Dotted line indicates chance level (50%). Statistical analysis calculated using a binomial test. **p<0.01; ***p<0.001.

**Figure S5 (related to Figure 3).**
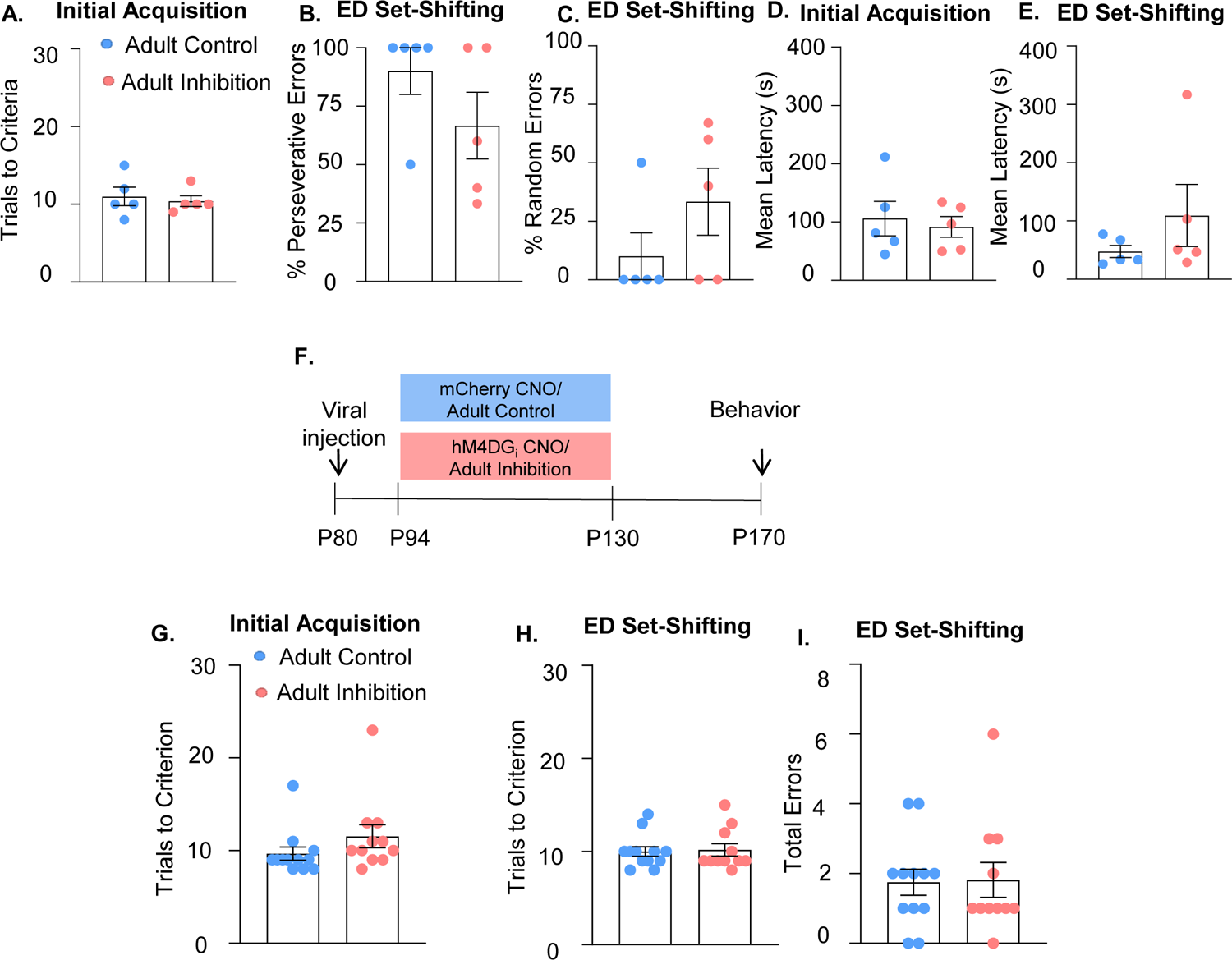
Additional behavioral analysis related to mPFC PV adult inhibition. **(A)** Trials to reach criterion in the initial acquisition phase of the set-shifting task is not affected by adult inhibition of mPFC PV interneurons. **(B-C)** There is no difference in the percentage of perseverative **(B)** or random **(C)** errors made by Adult Inhibition and Control mice. Adult Inhibition and Control animals have a comparable mean latency to make a choice in both the initial acquisition **(D)** as well as extradimensional phase **(E)** of the set-shifting task. **(F)** Schematic illustrating timeline for adult inhibition experiment where the virus is injected at P80 rather than P10. CNO was administered between P94 and P130 and behavior was assessed from P170 onwards. **(G)** Adult Inhibition animals with virus injected at P80 rather than P10 still take a comparable number of trials to reach criterion in both the initial acquisition as well as extradimensional phase **(H)** of the set-shifting task as Adult Controls. **(I)** They also make a comparable number of errors during ED set-shifting. In all graphs, dots show individual animal values and bars show the mean ± SEM. Statistical analysis performed with unpaired t-tests.

**Figure S6 (related to Figure 4).**
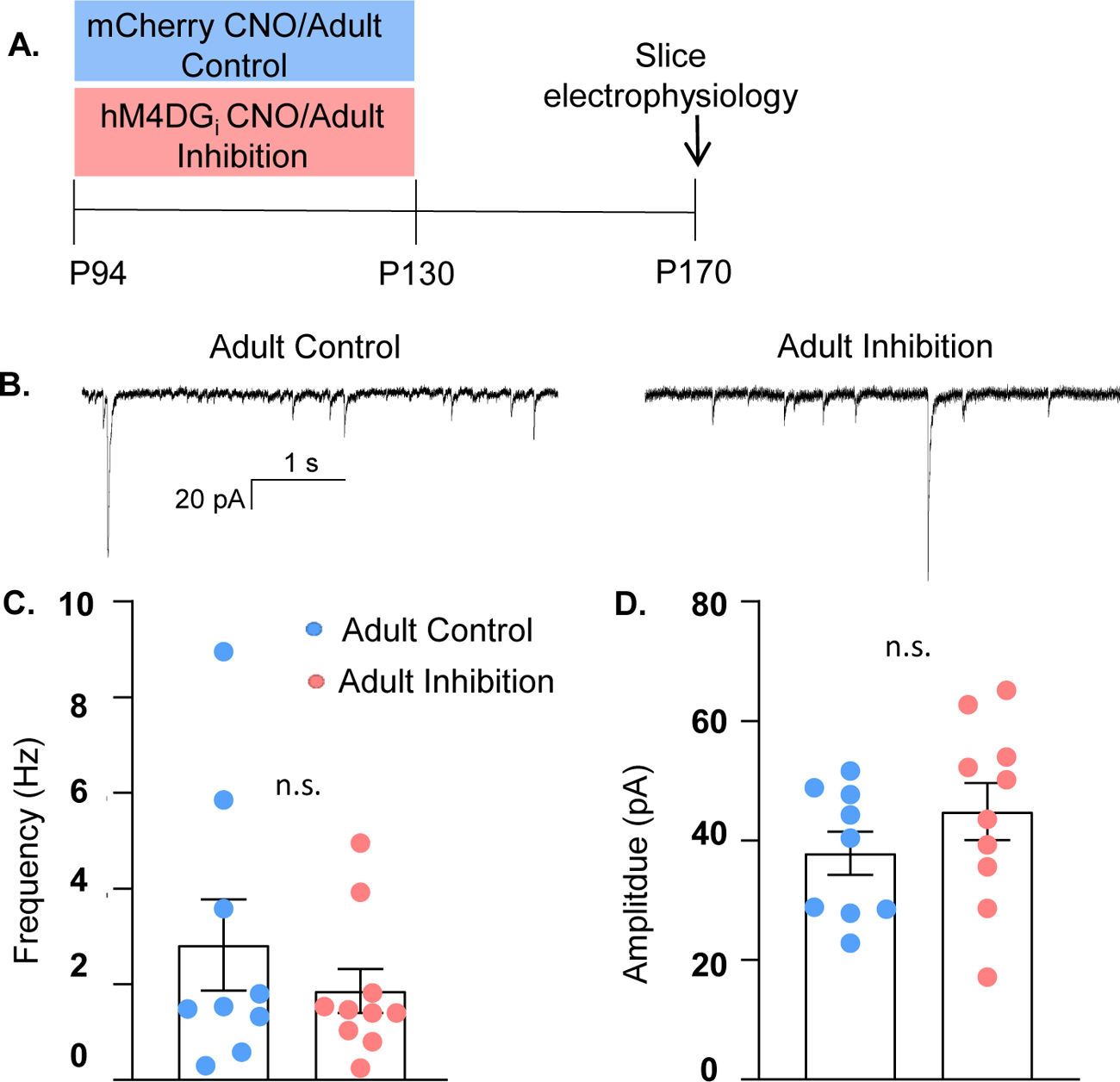
Adult mPFC PV inhibition does not lead to long-lasting effects on their functional inhibition of glutamatergic pyramidal cells. **(A)** Experimental schematic. PV-hM4DGi-mCherry or PV-mCherry expressing mice were injected with CNO from P94 to 130. Slice electrophysiological recordings of sIPSCs were made at P170. **(B)** Example current traces from Adult Control (left) and Adult Inhibition (right) mice. **(C)** There is no difference in the frequency or **(D)** amplitude of sIPSCs recorded from Adult Inhibition or Control animals. Statistical analysis performed with an unpaired t-test. n.s. = not significant.

**Figure S7 (related to Discussion).**
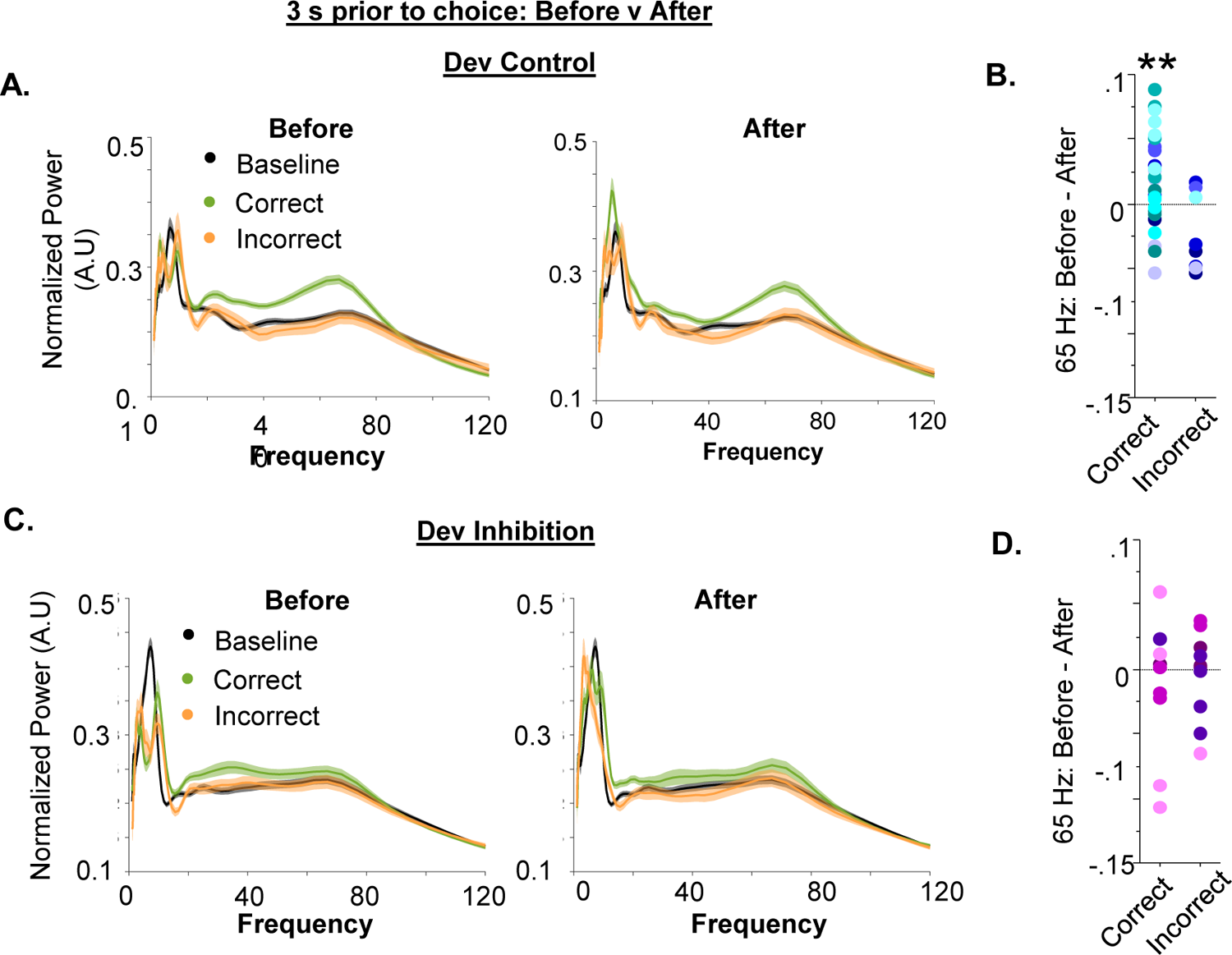
Additional analyses of gamma power during the set-shifting task. **(A)** Power as a function of frequency during the baseline (black) and in a 3-second window before (left) or after (right) choice in the set-shifting task, split by whether the choice was correct (green) or incorrect (orange) for Dev Control mice. **(B)** Quantification of the difference in 65 Hz frequency range power in the period before versus after a correct or incorrect choice. 65 Hz frequency range power was significantly greater before versus after making a correct choice (before-after mean+SEM: 0.02067±0.007246, n=31 trials from 8 Dev Control mice; **p=0.0078) but did not change for incorrect trials (before-after mean+SEM: −0.02337±0.01052, n=8 trials from 6 Dev Control mice; p=0.0618). **(C-D)** Same data as in (A-B) but for Dev Inhibition mice. 65 Hz frequency range power did not change before versus after making either a correct (before-after mean+SEM: −0.01359±0.01594, n=10 trials from 4 Dev Control mice; p=0.4160), or incorrect (before-after mean+SEM: −0.004430±0.01071, n=10 trials from 4 Dev Control mice; p=0.6889), choice Dots indicate data for individual trials color-coded by animal. Statistical analysis performed one-sample t-test compared to hypothetical mean of 0. **p<0.05.

